# Insights into the controls on metabolite distributions along a latitudinal transect of the western Atlantic Ocean

**DOI:** 10.1101/2021.03.09.434501

**Authors:** Winifred M. Johnson, Melissa C. Kido Soule, Krista Longnecker, Maya P. Bhatia, Steven J. Hallam, Michael W. Lomas, Elizabeth B. Kujawinski

## Abstract

Metabolites, or the small organic molecules that are synthesized by cells during metabolism, comprise a complex and dynamic pool of carbon in the ocean. They are an essential form of information, linking genotype to phenotype at the individual, population and community levels of biological organization. Characterizing metabolite distributions inside microbial cells and dissolved in seawater is essential to understanding the controls on their production and fate, as well as their roles in shaping marine microbial food webs. Here, we apply a targeted metabolomics method to quantify particulate and dissolved distributions of a suite of biologically relevant metabolites including vitamins, amino acids, nucleic acids, osmolytes, and intermediates in biosynthetic pathways along a latitudinal transect in the western Atlantic Ocean. We find that, in the euphotic zone, most particulate or intracellular metabolites positively co-vary with the most abundant microbial taxa. In contrast, dissolved metabolites exhibited greater variability with differences in distribution between ocean regions. Although fewer particulate metabolites were detected below the euphotic zone, molecules identified in the deep ocean may be linked to preservation of organic matter or adaptive physiological strategies of deep-sea microbes. Based on the identified metabolite distributions, we propose relationships between certain metabolites and microbial populations, and find that dissolved metabolite distributions are not directly related to their particulate abundances.

## Introduction

Marine organic matter is a complex component of the global carbon cycle due to its molecular diversity across particulate and dissolved phases. Cycling of this organic matter is largely controlled by the relationship between microbial metabolism and dissolved organic carbon (DOC) as marine microbes produce, consume, and influence the composition of DOC. About 50% of the organic matter fixed by phytoplankton passes through the marine DOC pool (Ducklow 1999), which is a carbon reservoir similar in size to that of carbon dioxide in the atmosphere (Hansell et al. 2009). Within the marine DOC pool, organic molecules are cycled on timescales ranging from minutes to thousands of years (reviewed by Carlson and Hansell 2015).

To adequately understand the factors that control production and removal of organic matter, molecular-level characterization of this material is needed. Compositional studies of organic matter have predominantly focused on compound classes. While a much smaller portion of the organic matter reservoir in the ocean, suspended particulate organic matter can be a source of dissolved organic matter through processes like cell death, grazing, and viral lysis (Carlson and Hansell 2015). These particles are composed of biologically-derived organic matter in the form of living cells and detritus including carbohydrates, amino acids, lipids, nucleic acids, and additional small molecules required for metabolism and secondary metabolites (e.g. Biersmith and Benner 1998; Sheridan et al. 2002; Kawasaki et al. 2011; Johnson et al. 2020).

Studies of molecular classes have determined that a large portion of high-molecular weight DOC is comprised of carbohydrates (Aluwihare et al. 1997) as well as carboxyl-rich aliphatic material (Hertkorn et al. 2006). Hydrolysable and free individual amino acids make up a smaller portion of organic carbon (McCarthy et al. 1996; Kaiser and Benner 2009). An additional group of molecules in DOC, identified with ultrahigh-resolution mass spectrometry of solid phase-extracted DOC, includes compounds with molecular formulas that do not match common biochemical compound classes, likely derived from pyro- and petrogenic sources (Dittmar and Koch 2006).

Metabolites, including free amino acids and monomeric sugars as well as vitamins and metabolic intermediates, while comprising a relative minor proportion of DOC in the ocean, play essential roles as the currencies in marine microbial interactions, transferring energy, cellular components, and information among microbial neighbors (Kujawinski 2011; Moran et al. 2016). By simultaneously measuring a suite of these molecules, we can trace the relationships between their distributions in particulate and dissolved phases, and hypothesize about the controls on their production, removal, and roles in microbial communities.

Metabolites, or small biomolecules, are defined as the products of all cellular regulation (Fiehn 2002). Due to the sensitivity of metabolite concentrations to both genetic and environmental factors, the metabolite profile of an organism can be considered an aspect of its phenotype. Intracellular or particulate metabolite concentrations respond to physical, chemical, and biological environmental cues including nutrient limitation (Brauer et al. 2006; Kujawinski et al. 2017), salinity (Gebser and Pohnert 2013), temperature (Thompson et al. 1992), oxidative stress (Lesser 2006), grazing (Pohnert 2000; Longnecker and Kujawinski 2020), viral lysis (Ankrah et al. 2014), available carbon substrates (Johnson et al. 2016), and the presence of infochemicals (Seyedsayamdost et al. 2011). For example, in response to infection by a phage, a marine bacterium had elevated intracellular concentrations of some amino acids and sugars (Ankrah et al. 2014). Thus, probing particulate and dissolved metabolite concentrations within the contexts of environmental cues and microbe-microbe interactions may elucidate phenotypic variations in microbes under environmental stressors, and thus facilitate understanding of exuded or released microbial organic matter under various conditions (Kujawinski 2011; Moran et al. 2016).

Dissolved metabolites mediate interactions among marine organisms. This encompasses both the transfer of organic substrates through the food web as well as the subtle but often significant role of infochemicals in influencing an organism’s phenotype (e.g. Paul et al. 2012; Barak-Gavish et al. 2018). Heterotrophic microbes catabolize many metabolite types produced by autotrophic microbes. For example, sugars fuel a dominant component of the marine food web, supporting aerobic respiration (Søndergaard et al. 2000). A common algal metabolite, dimethylsulfoniopropionate (DMSP), is used for both energy and reduced sulfur (Kiene et al. 2000). Amino acids also move through the food web, providing both fixed nitrogen and essential cellular building blocks (Rich et al. 1997). In addition to the autotrophic production of organic substrates for heterotrophic remineralization, some metabolites fulfill requirements in sympatric community members. For instance, certain species of phytoplankton require vitamins that they cannot synthesize *de novo*, such as vitamin B_12_ or biotin; these vitamins are produced by heterotrophic bacteria and sustain phytoplankton growth (reviewed by Croft et al. 2006). Some heterotrophic bacteria also have specific substrate requirements, such as a SAR11 isolate that requires an exogenous source of a thiamin precursor (Carini et al. 2014). These physiological demands have been studied on a case-by-case basis using field and lab experiments coupled with genomic information. However, our knowledge of the distributions of many of these metabolites in the ocean is limited or non-existent, thus inhibiting our ability to predict the impact of these requirements on microbial communities in the field.

To quantify a suite of structurally diverse metabolites, we use a liquid chromatography-tandem mass spectrometry-based metabolomics approach (Kido Soule et al. 2015). In this targeted metabolomics method, pure standards of each metabolite of interest are used to identify and quantify that metabolite in seawater. In this study, seawater samples encompassed a variety of ocean regions, between the latitudes of 55°N and 38°S in the western Atlantic and from depths between 5 m and ∼5500 m. Our method detected 27 particulate metabolites and 18 dissolved metabolites, including amino acids, vitamins, nucleic acids, osmolytes, and a variety of metabolic intermediates from both primary metabolism and biosynthetic pathways. Many of these metabolites were selected because they are essential for most organisms, while others were identified in laboratory experiments as being responsive to environmental differences or being specific to certain groups of organisms. Some of these metabolites have been measured previously in the ocean, allowing us to compare our measurements to literature values while other metabolites have never been measured, to our knowledge, in the ocean.

The goal of this study is to quantify the distribution of the core set of metabolites in our method along latitudinal and depth gradients in the Atlantic Ocean. In particular, the identification of metabolites whose distributions are independent of microbial biomass may lead to discovery of other environmental controls on the production or degradation of those molecules in certain ocean regions. As the majority of the metabolites analyzed are essential components of metabolism, starting from the null hypothesis that each metabolite is present in the same abundance inside a cell anywhere in the ocean and that its release into, and uptake from, the dissolved pool of metabolites is driven only by the number of cells present, particulate and dissolved metabolite distributions should co-vary with biomass. However, we find that this is not the case, and metabolite concentrations correlate to a variety of factors. We distill our results within the following three contexts: 1) particulate metabolite distributions that are correlated to abundant microbial community members, 2) dissolved metabolite distributions that vary across ocean regions and 3) particulate metabolite distributions in the euphotic zone versus the deep ocean.

## Materials and methods

### Materials

We obtained all metabolite standards from Sigma-Aldrich at the highest purity available with the exception of dimethylsulfoniopropionate (DMSP), which we purchased from Research Plus, Inc. We purchased hydrochloric acid (trace metal grade), acetonitrile (Optima grade), and methanol (Optima grade) from Fisher Scientific. We obtained formic acid (LC-MS grade) from Fluka Analytical. We purchased glutamic acid-d_3_ from Cambridge Isotopes, 4-hydroxybenzoic acid-d_4_ from CDN Isotopes, and sodium taurocholate-d_5_ from Toronto Research Chemicals through Fisher Scientific. We used water purified by a Milli-Q system (Millipore; resistivity 18.2 MΩ•cm @ 25 °C, TOC < 1 µM) for all cleaning, eluents, and solutions. We combusted all glassware and GF/F filters in an oven at 460°C for at least 5 h. We acid-washed and autoclaved all plasticware before use. We flushed the filter holders and tubing for shipboard filtration with 10% HCl and then Milli-Q water between each sample.

### Field Sites

We collected samples over the course of two cruises in 2013. Cruise KN210-04 (Deep DOM) took place in the western tropical Atlantic from 25 March – 9 May (austral fall), transiting from Montevideo, Uruguay to Bridgetown, Barbados. In the north Atlantic, we collected samples on the second leg of cruise AE1319 transiting from Boothbay Harbor, Maine U.S.A. to Bermuda via the Labrador Sea from 20 August – 11 September (boreal late summer – early fall). On KN210-04 we collected samples at depths including 5 m (referred to as the surface), the deep chlorophyll maximum (DCM) determined by fluorescence, 250 m, Antarctic Intermediate Water (AAIW, ∼1000 m), North Atlantic Deep Water (NADW, ∼2500 m), and Antarctic Bottom Water (AABW, ∼5000 m). On cruise AE1319 we collected samples at 5 m, the DCM, Eighteen Degree Mode Water (∼350 m, where present), ∼1000 m, and 3000 m. See Table S1 for mixed layer, euphotic zone, and DCM depths.

### Total organic carbon (TOC)

We collected 40 mL samples of unfiltered seawater in combusted glass EPA vials. We acidified the water samples to pH 2-3 with concentrated hydrochloric acid and stored them at 4°C until analysis. We analyzed the samples on a Shimadzu TOC-VCSH total organic carbon analyzer coupled to a TNM-1 analyzer. We ran Milli-Q water blanks and standard curves of potassium hydrogen phthalate and potassium nitrate throughout analysis and made comparisons daily to standards from Prof. D. Hansell (University of Miami).

### Temperature, salinity, photosynthetically active radiation (PAR), chlorophyll *a*, and cell counts

The R/V *Knorr* (KN210-04) was equipped with a SBE9+ CTD with a depth limit of 6000 m. We used a SBE3T/SBE4C sensor system to measure temperature and conductivity, a Wet Labs FLNTURTD combination fluorometer and turbidity sensor to detect fluorescence, and a Biospherical QSP-200L underwater PAR sensor. The R/V *Atlantic Explorer* (AE1319) had a SBE 9/11 Plus CTD with a depth limit of 6800 m with a SBE 3+ temperature sensor, a SBE 4C conductivity sensor, and a Biospherical QSP-2350 Scalar PAR sensor. We used a Chelsea Aquatracka II to measure fluorescence. We calibrated fluorescence data from both cruises with direct chlorophyll *a* measurements (according to methods from Arar & Collins (1997) on KN210-04; according to methods from Yentsch and Menzel (1963) on AE1319). Prokaryotic cell counts for cruise KN210-04 were based on epifluorescence microscopy of cells stained with SYBR Green I stain and captured on a 0.02 µm Anodisc filter (for a complete method see Noble and Fuhrman 1998).

### Shipboard sample processing

We collected water (4 L) directly from Niskin bottles into polytetrafluoroethylene (PTFE) or polycarbonate bottles. We filtered water sequentially through a 0.7-µm (nominal pore size) GF/F filter (Whatman) and a 0.2-µm filter (Omnipore, EMD Millipore) using a peristaltic pump. We re-wrapped the GF/F filters in their combusted aluminum foil envelopes and we folded and placed the Omnipore filters into cryogenic vials (Nalgene). We stored filters at −80°C until they could be extracted in the laboratory (next section).

We then acidified the filtrate with 4 mL of 12 M HCl (∼pH 2-3; Dittmar et al. 2008; Longnecker 2015). We extracted dissolved organic molecules from the filtrate using solid phase extraction (SPE) with a modified styrene-divinylbenzene polymer (Agilent Bond Elut PPL) as a substrate. We first rinsed the PPL cartridges with 6 mL of methanol and then used a vacuum pump to pull the acidified filtrate (4 mL of 12 M HCl added to 4 L of filtrate (∼pH 2-3) through the cartridge via PTFE tubing (Dittmar et al. 2008; Longnecker 2015). Then we rinsed the cartridge with ∼24 mL of 0.01 M HCl and allowed it to dry by pulling air over the cartridge for 5 min. We eluted each sample with 6 mL of methanol into a glass test tube and then transferred it via Pasteur pipette to an 8 mL amber vial. We stored extracts at −20°C until analysis. We created process blanks by carrying shipboard and laboratory Milli-Q water through the entire procedure of filtration and SPE extraction.

### Laboratory sample processing

We extracted particulate filters within 48 h of mass spectrometry analysis. We adapted the filter extraction protocol (Kido Soule et al. 2015) from Rabinowitz and Kimball (2007). We weighed one half of each filter and cut it into smaller pieces using methanol-rinsed scissors and tweezers on combusted aluminum foil. We placed the pieces in an 8 mL glass amber vial with 1 mL of −20°C extraction solvent (40:40:20 acetonitrile:methanol:water + 0.1 M formic acid) and spiked 25 µL of a 1 µg/mL deuterated standard mix (glutamic acid-d_3_, 4-hydroxybenzoic acid-d_4_, taurocholate-d_5_) into each sample as an extraction recovery standard. We vortexed each vial gently to separate filter pieces and then sonicated the vials in an ultrasonication bath for 10 min. We transferred the solvent extract with a Pasteur pipette to 1.5 mL Eppendorf centrifuge tubes. We rinsed the filter pieces left in the 8 mL vials with 200 µL of cold extraction solvent and combined the rinse with the original extract. We spun the extract at 20,000 x *g* for 5 min to remove cellular detritus and filter particles, and transferred the supernatant to clean 8 mL amber glass vials for neutralization with 26 µL of 6 M ammonium hydroxide. We removed the solvent by vacufuge until samples were almost completely dry (<5 µL) and reconstituted the dry extract with either 247.5 µL 95:5 water:acetonitrile and 2.5 µL biotin-d_4_ (injection standard) for samples collected at the surface or DCM, or 123.75 µL 95:5 water:acetonitrile and 1.25 µL biotin-d_4_ (injection standard) for deeper samples with presumed lower biomass. We loaded 100 µL of each solution into a glass insert in an autosampler vial. We combined 15 µL of each sample for matrix-matched quality control during the LC-MS/MS analysis (see next section).

For the dissolved metabolites, we brought 1 mL of the SPE extracts (∼6 mL total volume) to almost complete dryness in the vacufuge and then reconstituted the samples in 495 µL 95:5 water:acetonitrile and 5 µL of 5 µg/mL biotin-d_4_ (injection standard). Some dissolved metabolite samples (mostly deep samples, 2000 – 5000 m) required additional dilution due to high-levels of ion suppression in the middle of the chromatogram (see Table S2).

**Table 2.** Range of dissolved concentrations of each metabolite and prevalence in samples. Minimum concentration is the lowest value in a sample where the metabolite was detected. Values with a “<” symbol indicate the detection limit of the metabolite as the metabolite was not detected in any sample.

| Compound | Abbreviation | Min<br>(pM)<br>(>0) | Max<br>(pM) | # of samples<br><200m<br>(n=26) | # of samples<br>>200m<br>(n=28) |
| --- | --- | --- | --- | --- | --- |
| 2,3-Dihydroxybenzoic acid | 2,3-DHBA | 4.2 | 9.9 | 3 | 0 |
| 3-Mercaptopropionic acid |  | < 78 | < 78 | 0 | 0 |
| 4-Aminobenzoic acid | PABA | 16 | 51 | 25 | 0 |
| 4-Hydroxybenzoic acid | 4-HBA | 15 | 61 | 7 | 0 |
| 5-Methylthioadenosine | MTA | 1.1 | 2.6 | 9 | 0 |
| Adenosine |  | < 123 | < 123 | 0 | 0 |
| Biotin |  | 3.5 | 26 | 6 | 2 |
| Caffeine |  | 2.6 | 8.7 | 12 | 4 |
| Chitotriose |  | 12 | 130 | 7 | 1 |
| Cyanocobalamin |  | < 0.8 | < 0.8 | 0 | 0 |
| Desthiobiotin |  | 27 | 27 | 1 | 0 |
| Folic acid |  | < 0.7 | < 0.7 | 0 | 0 |
| Indole 3-acetic acid |  | 5.7 | 5.7 | 1 | 0 |
| Inosine |  | 16 | 29 | 2 | 0 |
| (Iso)leucine |  | 90 | 230 | 4 | 0 |
| *N*-Acetylglutamic acid |  | 220 | 600 | 1 | 1 |
| Nicotinamide adenine<br>dinucleotide | NAD | 56 | 56 | 1 | 0 |
| Pantothenic acid |  | 1.0 | 19 | 22 | 0 |
| Phenylalanine |  | 2.2 | 290 | 26 | 12 |
| Pyridoxine |  | < 236 | < 236 | 0 | 0 |
| Riboflavin |  | 0.7 | 12 | 10 | 7 |
| *S*-(5'-Adenosyl)-L-homocysteine |  | < 72 | < 72 | 0 | 0 |
| Taurocholic acid |  | < 5 | < 5 | 0 | 0 |
| Thiamin |  | 4000 | 4000 | 1 | 0 |
| Thymidine |  | < 123 | < 123 | 0 | 0 |
| Tryptophan |  | 20 | 190 | 18 | 1 |

### Mass spectrometry

We performed the LC-MS/MS analysis on a reversed phase C18 column (Phenomenex Synergi Fusion, 2.1 x 150 mm, 4 µm) coupled via heated electrospray ionization (ESI) to a triple quadrupole mass spectrometer (Thermo Scientific TSQ Vantage) operated under selected reaction monitoring mode (SRM), as described previously (Kido Soule et al. 2015). We monitored quantification and confirmation SRM transitions for each analyte. Chromatography conditions included a gradient between Eluent A (Milli-Q water with 0.1% formic acid) and Eluent B (acetonitrile with 0.1% formic acid) at 250 µL / min: hold at 5% B for 2 min; ramp to 65% B for 16 min; ramp to 100% B for 7 min and hold for 8 min. We re-equilibrated the column with the starting ratio of eluents for 8.5 min between samples. We separated samples into batches of approximately 50 samples, with a pooled sample comprised of those 50 samples. We conditioned the column with 5 injections of the pooled sample prior to each batch, and we ran a pooled QC sample after every ten samples. These samples indicated that coefficients of variation of metabolite abundance ranged between 8-27% during mass spectrometry analysis.

### Data processing

We converted the XCalibur RAW files generated by the mass spectrometer to mzML files using msConvert (Chambers et al. 2012). We used MAVEN (Melamud et al. 2010; Clasquin et al. 2012) to select and integrate peaks. We discarded calibration peaks below a MAVEN quality threshold of 0.4 (on a scale of 0-1) and sample peaks below 0.2. To enhance confidence in metabolite identification, we required quantification and confirmation peaks to have retention times within 12 seconds (0.2 minutes) of each other. We required confirmation ions to have a MAVEN quality score of at least 0.1 and a signal-to-noise ratio greater than 1. We required calibration curves to have at least five calibration points, with the highest point at one concentration level above the highest concentration in a sample. There were 9 available calibration points: 0.5 ng mL^−1^, 1 ng mL^−1^, 10 ng mL^−1^, 25 ng mL^−1^, 50 ng mL^−1^, 100 ng mL^−1^, 250 ng mL^−1^, 500 ng mL^−1^, 1000 ng mL^−1^. We adjusted particulate metabolite abundances based on ion suppression determined by comparison of a standard spiked pooled sample (1000 ng mL^−1^ metabolite standard mix in the sample matrix) to a standard spiked Milli-Q water sample (1000 ng mL^−1^ metabolite standard mix without complex matrix), (see Johnson et al. 2020 for ion suppression values). We normalized metabolite abundances to the volume of seawater filtered. Where noted, we normalized particulate concentrations of metabolites to the total moles of targeted metabolites measured in the sample because this parameter correlated well with available cell counts (see discussion below and Figure S1). We corrected the concentrations of dissolved metabolites with extraction efficiencies greater than 1% on a PPL cartridge (measured concentration divided by the extraction efficiency) to reflect a more accurate estimate of the *in situ* metabolite concentrations (see extraction efficiencies in Johnson et al. 2017). We do not report dissolved metabolites with extraction efficiencies less than 1%.

### Determination of microbial community composition

We collected different types of microbial community composition data on the southern transect (KN210-04) and the northern transect (AE1319). In the south Atlantic we used small subunit ribosomal RNA (16S or SSU rRNA) gene amplicon sequencing to determine bacterial and archaeal community composition as described in Johnson et al. (2020). In the north Atlantic, we used flow cytometry methods to determine abundances of *Synechococcus*, *Prochlorococcus*, picoeukaryotes, and nanoeukaryotes as described in Lomas et al. (2014).

### Computation and statistical tools

We used MATLAB (R2014a; MathWorks, Natick, MA) to process data and create figures. We created the map and metabolite profile images with Ocean Data View (Schlitzer 2016). We calculated Spearman’s rank correlation coefficients using the R stats package (R Core Team 2015) to examine the relationship between metabolite distributions and microbial community composition. Spearman’s rank correlation identifies monotonic relationships but does not require that the relationship be linear. We required that metabolites be present in at least three samples to perform the analysis. When working with the phylogenetic data from cruise KN210-04 we only included the top ten most abundant phyla. We adjusted the p-values from the correlation analysis using the false discovery rate algorithm in the R package q-value (Storey 2002, 2010; Storey and Tibshirani 2003). We considered the resulting adjusted q-values significant at the value permitting the possibility of one false positive for the smaller north Atlantic dataset (*p* < 0.06 particulate metabolites; *p* < 0.35 dissolved metabolites). For the larger south Atlantic dataset we defined the significance cut-off as *p* < 0.05. We defined mixed layer depth as the depth where the decreasing temperature exceeded 0.15 °C m^−1^ (based upon the continuous CTD downcast) and the euphotic zone depth as the depth at which photosynthetically active radiation (PAR) was 1% of the surface PAR.

## Results

### Oceanographic transect

We collected samples along a transect encompassing latitudes from 38°S to 55°N and at depths ranging from the surface (5 m) to approximately 5500 m. Surface oceanic regions along this transect included the South Atlantic Gyre (Stations K2, K5, K7, K9), the equatorial region (Stations K12, K15, K16, K19, K21), the Amazon River plume (Station K23), the North Atlantic Gyre (Stations A11, A15), and the Labrador Sea (Stations A4, A7; Figure 1a) facilitating an analysis of metabolite distributions in both productive and oligotrophic regions of the ocean. In addition, we took samples from deep water masses including Antarctic Intermediate Water, North Atlantic Deep Water, and Antarctic Bottom Water (Figure S2). Stations and depths with likely high productivity were identified using chlorophyll *a* concentrations (Figure 1c). These included the deep chlorophyll maximum (DCM) near the equator where there was upwelling of nutrients (station K15) and the highest latitudes sampled in the Labrador Sea (stations A4, A7) where there was a shallow DCM. The total organic carbon (TOC) concentrations, as expected, were maximal in the surface water of the subtropical gyres and decreased with depth (Figure 1b).

**Figure 1.**
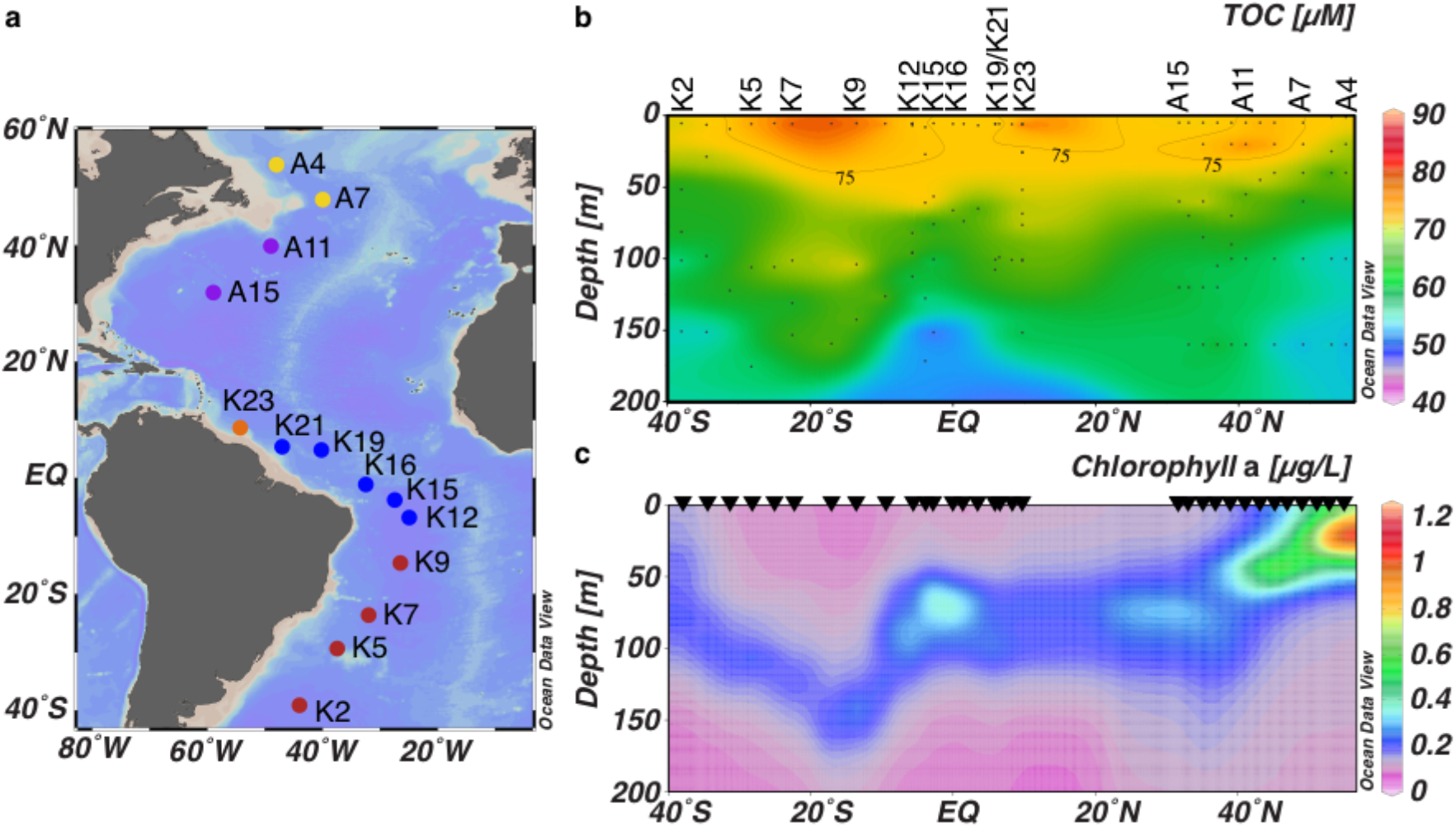
Transect stations and chemical parameters. (a) Stations sampled along a latitudinal transect in the western Atlantic Ocean. Dots colored by region: red, South Atlantic Gyre; blue, equatorial region; orange, Amazon plume; purple, North Atlantic Gyre; yellow, Labrador Sea. (b) The total organic carbon (TOC) concentration in the top 200 m along the transect. Black dots indicate sample locations. (c) Chlorophyll *a* concentrations in the top 200 m along the transect. Triangles indicate locations of CTD casts where fluorescence was measured.

### Data normalization

We report both particulate and dissolved metabolite concentrations per total volume filtered (picomoles per liter, pM). This approach is the most unambiguous way to present the data as concentration units are the standard chemical and geochemical approach to reporting chemical data. Concentration units also facilitate comparisons between the particulate and dissolved distributions. However, as particulate metabolite samples are approximations of the intracellular pool of metabolites, an approach that presents metabolite abundances in relation to cellular biomass is also needed. Other ‘omics fields use approaches such as normalization to housekeeping genes (whose abundances are constant inside cells) to present per-cell variability. Such an approach is not possible here because there is no consensus “housekeeping” metabolite. We considered other oceanographic parameters as proxies for biomass such as chlorophyll *a* concentration or the beam transmittance at 660 nm (from a transmissometer) as a measure of how many particles were in the water. Chlorophyll *a* is problematic because it is only produced by phytoplankton, and its abundance can be affected by physiological changes (Kruskopf and Flynn 2006). The relationship between transmissometer data and prokaryotic cell counts (from SYBR Green staining and counting under an epifluorescence microscope) showed a poor relationship. In this study, we predominantly use pM units, but present some data as the mole fraction where the moles of the metabolite are normalized to the total moles of metabolites measured in the sample. This was selected because there is good agreement between prokaryotic cell abundance and total moles of metabolites measured (Figure S1). As more oceanographic measurements of metabolites become available, best practices in normalization will inevitably emerge.

### Variability in metabolite detection

Of the 32 metabolites that passed the quality control checks of their calibration curves in the particulate samples, 5 were not detected in any sample. 2,3-Dihydroxybenzoic acid (2,3-DHBA), desthiobiotin, indole 3-acetic acid, and inosine 5′-monophosphate (IMP) were each measured in only one sample preventing assessment of their variability across the full transect. Most of the remaining metabolites were elevated in the upper 200 m of the water column, with DMSP, guanine, and phenylalanine measured in every particulate sample in this depth range. Throughout the full water column, guanine and phenylalanine were measured in all but 3 samples (Table 1).

**Table 1.** Range of particulate concentrations of each metabolite and prevalence in samples. Minimum concentration is the lowest value in a sample where the metabolite was detected. ^a^Picomoles of particulate metabolite per liter filtered. Values with a “<” symbol indicate the detection limit of the metabolite as the metabolite was not detected in any sample. ^b^Isoleucine and leucine cannot be separated chromatographically and thus their combined concentrations are reported as (iso)leucine.

| Compound | Abbreviation | Min<sup>a</sup> pM<br>(>0) | Max<sup>a</sup> pM | # of samples<br><200m<br>(n=34) | # of samples<br>>200m<br>(n=46) |
| --- | --- | --- | --- | --- | --- |
| 2,3-Dihydroxybenzoic acid | 2,3-DHBA | 1.6 | 1.6 | 1 | 0 |
| 3-Mercaptopropionate |  | < 12 | < 12 | 0 | 0 |
| 4-Aminobenzoic acid | PABA | < 0.6 | < 0.6 | 0 | 0 |
| 4-Hydroxybenzoic acid | 4-HBA | 0.9 | 4.4 | 2 | 2 |
| 5-Methylthioadenosine | MTA | 0.1 | 1.0 | 29 | 2 |
| Adenine |  | 0.6 | 22 | 11 | 21 |
| Adenosine |  | 0.09 | 85 | 29 | 17 |
| Adenosine 5'-monophosphate | AMP | 0.07 | 98 | 30 | 25 |
| Arginine |  | 2.4 | 46 | 9 | 22 |
| Biotin (vitamin B<sub>7</sub>) |  | 0.03 | 1.0 | 4 | 5 |
| Caffeine |  | 0.02 | 4.2 | 12 | 34 |
| Cytosine |  | < 2.2 | <2.2 | 0 | 0 |
| Desthiobiotin |  | 0.3 | 0.3 | 1 | 0 |
| Dimethylsulfoniopropionate | DMSP | 13 | 6200 | 34 | 8 |
| Folic acid |  | 0.2 | 0.8 | 2 | 2 |
| Glutamic acid |  | 61 | 9800 | 28 | 7 |
| Guanine |  | 1.3 | 1000 | 34 | 43 |
| Glycine betaine |  | 8.1 | 1300 | 33 | 7 |
| Indole 3-acetic acid |  | 0.2 | 0.2 | 1 | 0 |
| Inosine 5'-monophosphate | IMP | 0.9 | 0.9 | 1 | 0 |
| Inosine |  | 0.04 | 18 | 27 | 14 |
| (Iso)leucine<sup>b</sup> |  | 0.3 | 41 | 22 | 20 |
| Methionine |  | 1.5 | 39 | 4 | 2 |
| N-acetyl glucosamine |  | < 9 | < 9 | 0 | 0 |
| Pantothenic acid (vitamin B<sub>5</sub>) |  | < 6.7 | <6.7 | 0 | 0 |
| Phenylalanine |  | 0.05 | 58 | 34 | 43 |
| Proline |  | 0.2 | 110 | 23 | 25 |
| Pyridoxine (vitamin B<sub>6</sub>) |  | 0.1 | 2 | 8 | 20 |
| Riboflavin (vitamin B<sub>2</sub>) |  | 0.003 | 0.7 | 13 | 1 |
| Tryptophan |  | 0.02 | 20 | 33 | 32 |
| Uracil |  | 7.4 | 400 | 8 | 5 |
| Xanthine |  | 0.2 | 29 | 21 | 5 |

Twenty-six dissolved metabolites passed quality control criteria for their calibration curves and could be extracted using our PPL SPE method (Johnson et al. 2017). Of these, 8 were not detected in any samples. In some cases, this may be due to extremely low solid phase extraction efficiencies (i.e., high detection limits). No metabolites were measured in every sample in the dissolved phase. However, phenylalanine was the most prevalent of the dissolved metabolites, detected in 70% of these samples. 4-Aminobenzoic acid (PABA), pantothenic acid (vitamin B_5_), and tryptophan were present in most samples from the upper 200 m (Table 2). We examined the relationship between dissolved and particulate metabolites and based on the observed lack of a consistent relationship, we believe that filtration-induced cell leakage did not have a significant impact on our data and its interpretation (Text S1).

### Microbial taxa abundance

The datasets used to describe the composition of the microbial community were generated with different analytical approaches between the two cruises, and are not directly comparable. For both cruises, flow cytometry provided an assessment of the pigmented phytoplankton, including cyanobacteria *Prochlorococcus* and *Synechococcus*, picoeukaryotes and nanoeukaryotes. In the northern Atlantic (Figure S3), *Prochlorococcus* was numerically the largest of these four groups in the subtropics (Figure S3; stations A11 and A15) and a small component of stations A4 and A7, where *Synechococcus* and picoeukaryotes were most abundant (Baer et al. 2017). *Prochlorococcus* was numerically more abundant in the surface waters of the south and equatorial Atlantic than *Synechococcus*, picoeukaryotes, and nanoeukaryotes (Howard et al. 2017). In the southern and equatorial Atlantic, phytoplankton data were complemented by the prokaryotic community composition based on 16S rRNA gene sequences (Figure S4). Consistent with the literature on pelagic prokaryotic communities (Frias-Lopez et al. 2008), Cyanobacteria, Proteobacteria, and Bacteroidetes were the most abundant phyla in the upper 200 m (Figure S4). Cyanobacteria, specifically *Prochlorococcus*, was a more abundant component of the microbial community at the surface, at every station along this transect except for stations A4 and A7. This is consistent with the overall understanding of the distribution of these groups in the ocean (Zubkov et al. 1998; Li and Harrison 2001; Flombaum et al. 2013).

### Trends in particulate metabolites

Most particulate metabolites were elevated in the upper 200 m of the water column, relative to deeper samples (e.g. Figure 2) and 15 of the 27 quantified metabolites were found more frequently in the upper 200 m than below that depth (Table 1). These results are consistent with expectations based on the higher productivity and more abundant biomass in this zone. For example, guanine, a purine nucleobase required by all living organisms, was measured in nearly all the samples in the transect allowing comparison of its concentrations at all depths. Particulate guanine abundance was relatively high in the surface samples, generally peaking at the DCM, and then was quite low at depth (Figure 2a). As an essential component of DNA, guanine is a metabolite that might be hypothesized to co-vary with cell abundance. While the relationship between guanine and proxies of biomass such as prokaryotic cell abundance (r^2^ = 0.61, *p* < 0.001) and chlorophyll *a* concentration (r^2^ = 0.62, *p* < 0.001) was statistically significant, more study is required before it could be used as a proxy of biomass. When guanine is normalized to the total moles of targeted metabolites measured in a sample, it becomes a major component of the deep samples reaching greater than 70% of the total moles of metabolites in some samples (Figure 2b).

**Figure 2.**
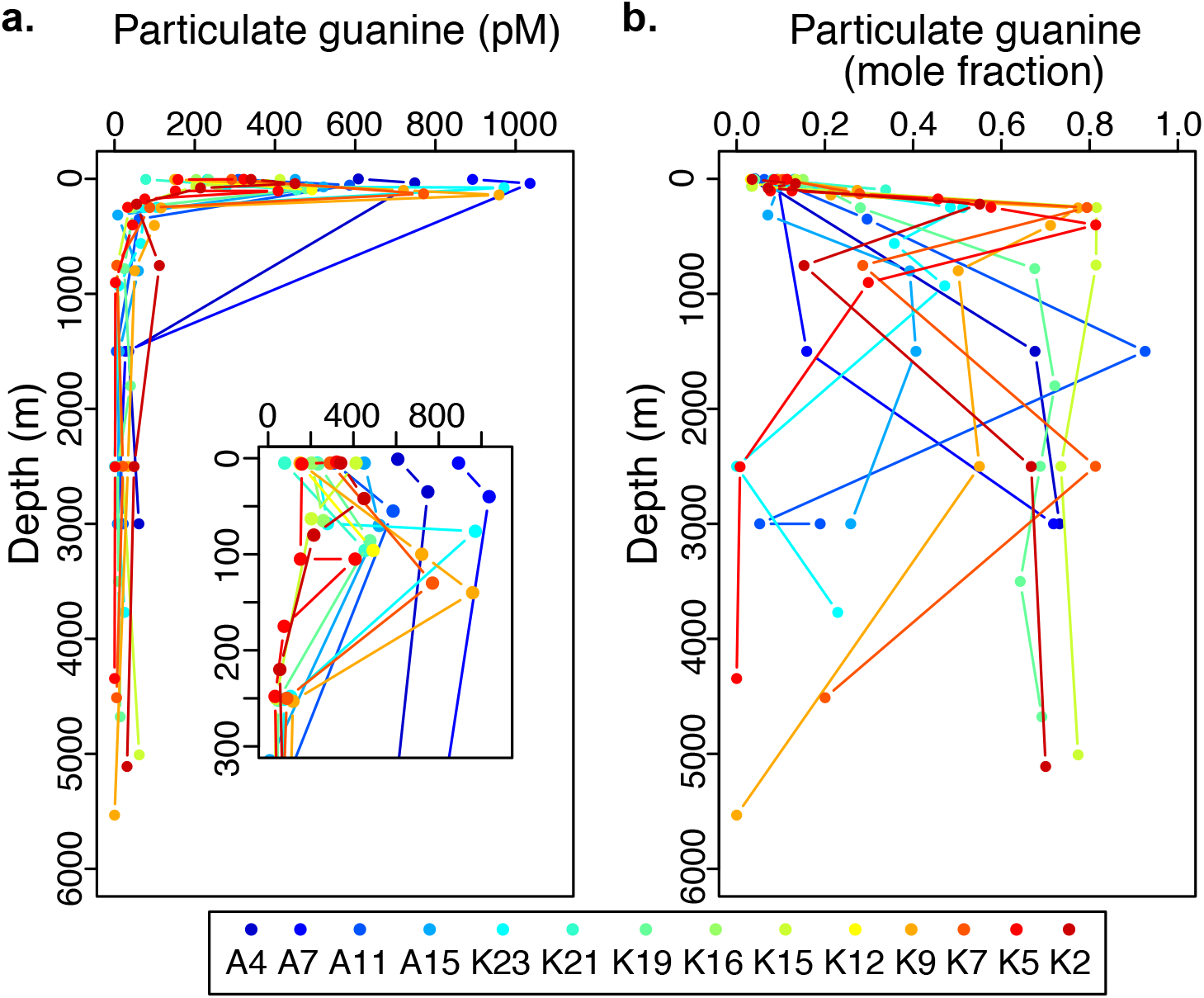
Particulate guanine abundance with depth. (a) Picomoles particulate guanine normalized to liter of seawater sampled, profiled by depth for each station. Cut-out shows the top 300 m of the water column in more detail. (b) Moles of guanine normalized to total moles targeted metabolites, profiled by depth for each station. Stations are indicated by different symbol colors, as shown in the legend.

To examine relationships between particulate metabolites and the distribution of microbial taxa, we used Spearman’s rank correlation coefficients to determine the extent of positive or negative co-variance. In the south and equatorial Atlantic, Cyanobacteria and Actinobacteria had the most significant positive correlations with many of the particulate metabolites, closely followed by Bacteroidetes and Proteobacteria (Figure 3a). Arginine and caffeine had significant positive correlations with other phyla (Figure 3a). In the north Atlantic, we observed a similar trend, wherein nanoeukaryotes, picoeukaryotes, and *Synechococcus* were generally positively correlated with most metabolites and *Prochlorococcus* was negatively correlated with metabolite abundance (Figure 3b). Uracil and adenine were the metabolites most weakly correlated with microbial taxa in both the south and north Atlantic. Additionally, in the south Atlantic biotin, folic acid, and methionine were not significantly correlated with any of the 10 most abundant phyla (Figure 3) suggesting that other factors control their distributions. Thus, particulate metabolites fall generally into two groups: 1) those that positively correlate with abundant taxa or 2) those that do not have correlations with specific taxa.

**Figure 3.**
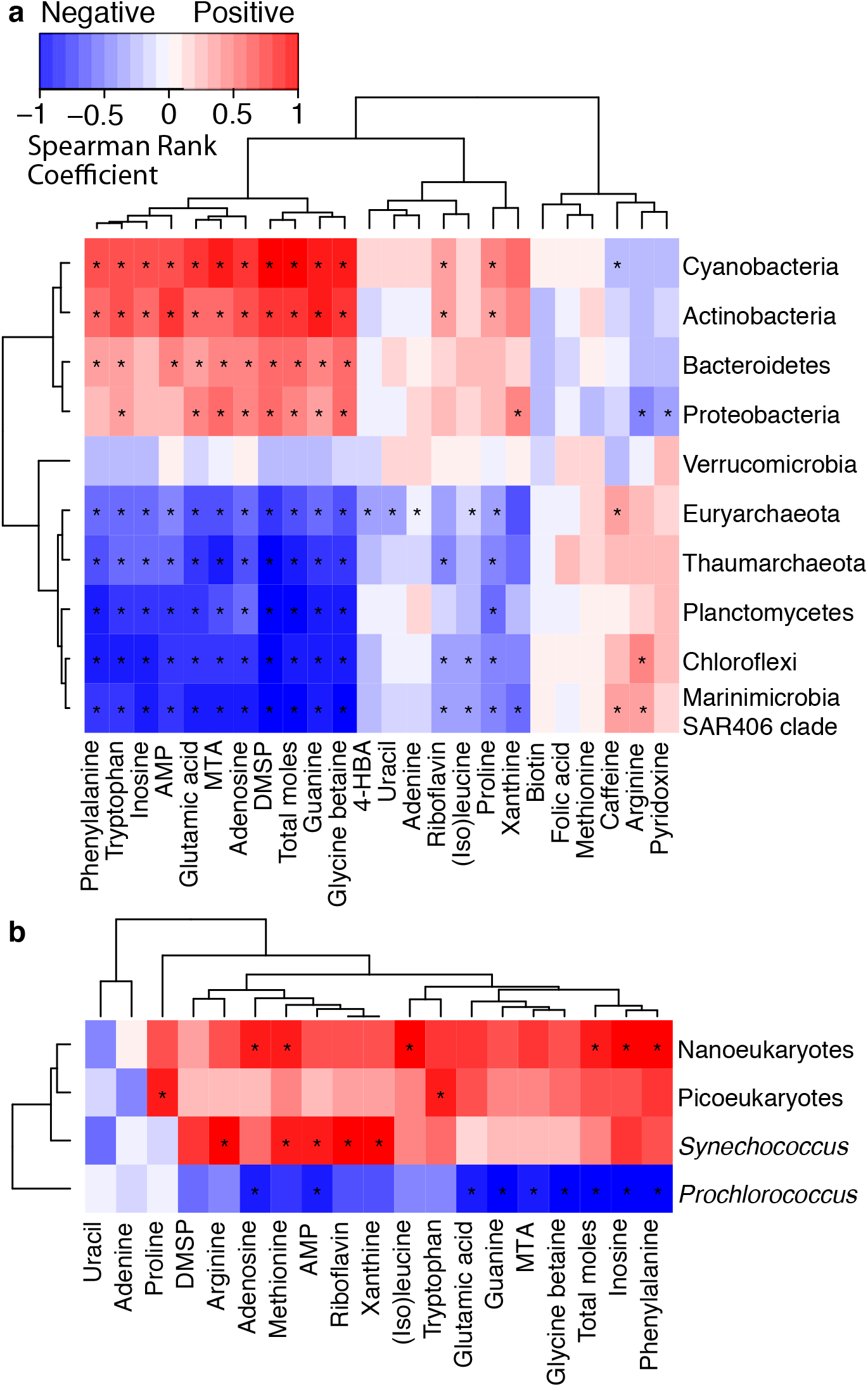
Spearman rank correlation coefficients plotted as a heatmap with hierarchical clustering showing the relationship between particulate metabolites and abundance of microbial community members. Red indicates a positive correlation and blue indicates a negative correlation. * Indicates correlations with significant *p*-values. (a) Comparison of metabolites to phyla from KN210-04 samples (*p* < 0.05). (b) Comparison of AE1319 metabolite samples from the surface and DCM to groups identified with flow cytometry (*p* < 0.06, see text).

Adenosine 5′-monophosphate (AMP) is representative of the distributions seen for metabolites that positively correlated with abundant taxa (Figure 4a) and had a linear (r^2^ = 0.79, *p* < 0.001) correlation with chlorophyll *a* concentrations (Figure 4b). These metabolites (phenylalanine, tryptophan, inosine, AMP, glutamic acid, 5-methylthioadenosine (MTA), adenosine, DMSP, guanine, glycine betaine) were characterized by elevated concentrations at high latitudes in the north Atlantic, at the DCM at K15 where upwelling was observed, and at the surface of K23 where there was some influence from the Amazon River plume. Relatively low concentrations were measured in the surface at K7, K9, and K12 in the South Atlantic Subtropical Gyre. These metabolites also grouped with the summed total moles of metabolites measured in each sample, indicating that they account for the bulk of the molar abundance of metabolites detected (Figure 3).

**Figure 4.**
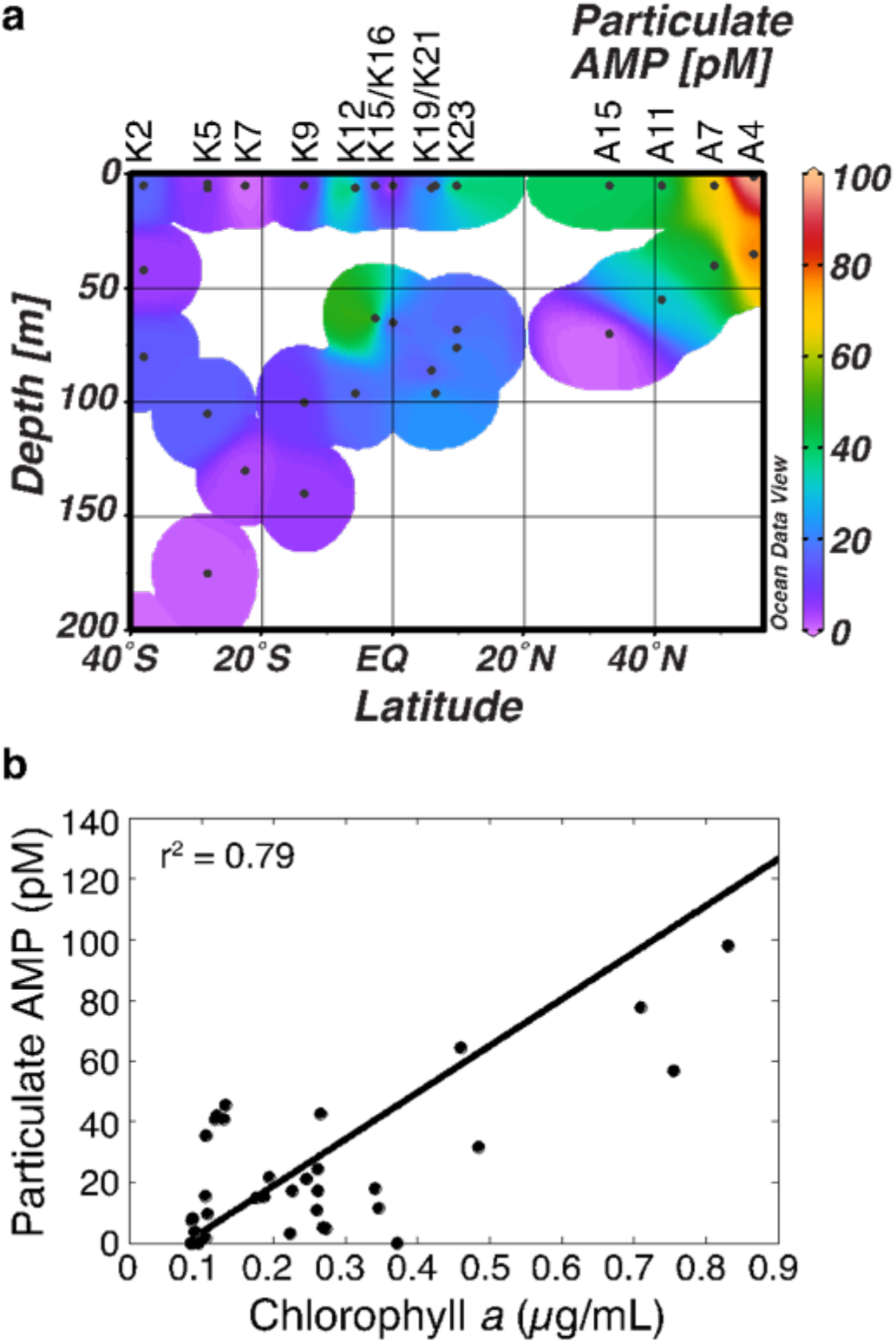
Particulate adenosine 5′-monophosphate (AMP) abundance with depth (a) and relative to chlorophyll *a* concentrations (b). (a) AMP abundance (pM) above 200 m. (b) Linear regression of chlorophyll *a* versus AMP, with r^2^ value 0.79 and *p* < 0.001 (slope = 154, y-intercept = −12).

Below 200 m, the amino acids phenylalanine, tryptophan, (iso)leucine, proline (Figure 5b), and arginine (Figure 5a), and the purine nucleobases guanine and adenine were detected (Table 1). Guanine in particular was found in most deep samples (Figure 2). Caffeine was also measured in deep samples and was strongly positively correlated with some of the other deep metabolites (guanine, phenylalanine, tryptophan). Given our limited knowledge of caffeine in the ocean, we cannot constrain whether caffeine’s presence is due to production by an organism as a defense compound similar to terrestrial plants (Mithöfer and Boland 2012) or from some other source, but its depth profile is not consistent with ship contamination. Finally, the B vitamins biotin (B_7_) and pyridoxine (B_6_), were measured below the euphotic zone. Biotin was measured in only a few deep samples while pyridoxine, an essential cofactor in many enzymes associated with amino acid metabolism (Mittenhuber 2001), was more prevalent. It is striking that of all the B vitamins measured, pyridoxine was present at a variety of depths with relatively consistent concentrations, suggesting that this vitamin might be of particular importance to deep sea microbes.

**Figure 5.**
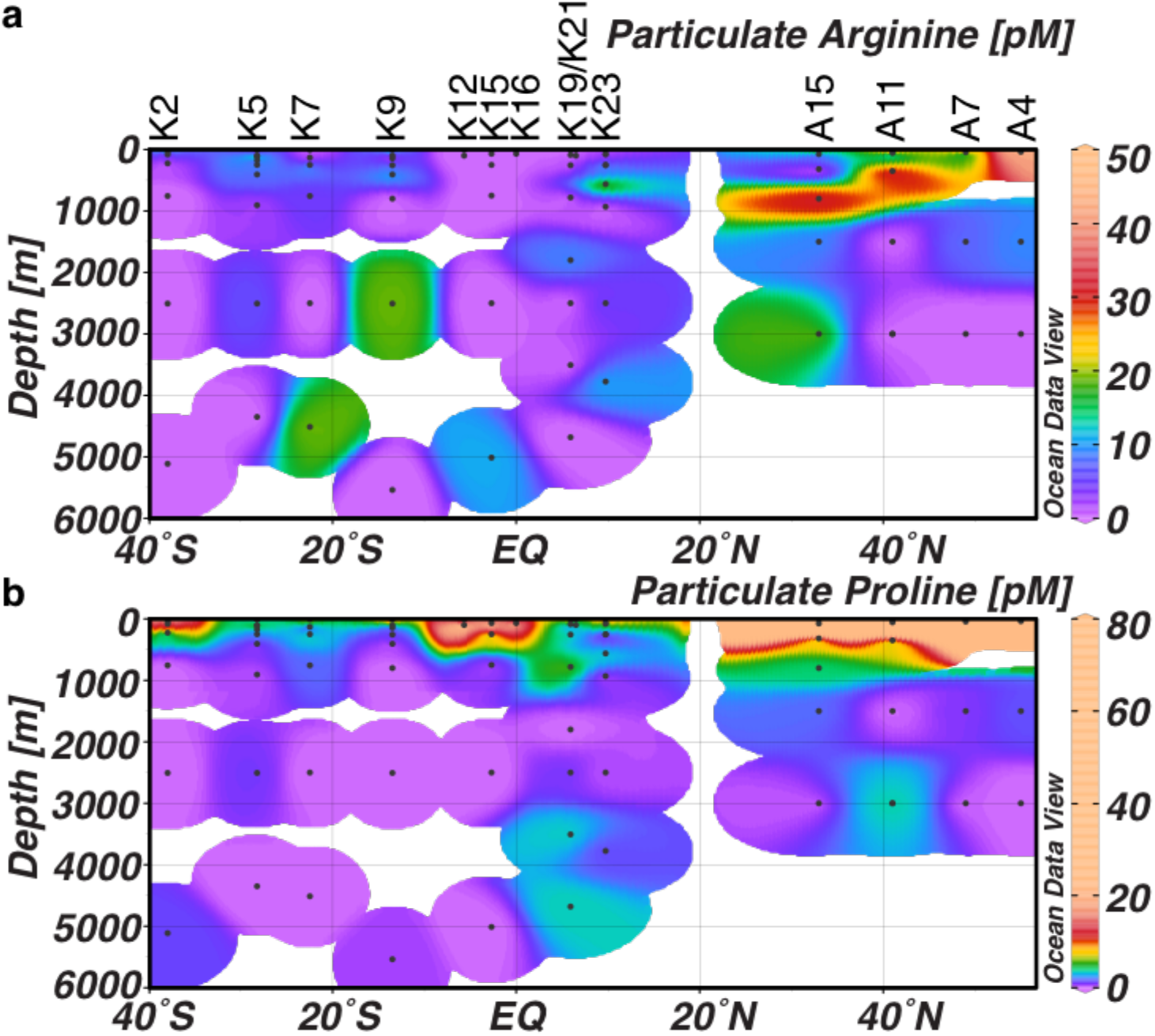
Distributions of particulate (a) arginine and (b) proline measured at depth.

### Trends in dissolved metabolites

The distributions of dissolved metabolites in the upper 250 m were weakly correlated with prokaryotic taxa in the south Atlantic. This correlation was conducted on all samples down to 250 m to provide enough statistical power to make the comparison and to include data that contrasted with the more productive surface and DCM samples. In the north Atlantic, the dissolved metabolites were positively correlated with either nanoeukaryotes and picoeukaryotes or with *Synechococcus* (Figure 6). While few correlations were significant after a false discovery rate correction, we observed similar trends within the correlations from both the south and north Atlantic, underscoring the potential links between microbial taxa and dissolved metabolites. For example, in the south Atlantic, dissolved PABA and pantothenic acid, had significant positive correlations with Cyanobacteria, likely driven by the abundance of *Prochlorococcus* (Figure 6a). In the north Atlantic, dissolved MTA was positively correlated with both picoeukaryotes and nanoeukaryotes, while pantothenic acid was positively correlated with *Synechococcus.* In the same region, *Prochlorococcus* was generally negatively correlated with the dissolved metabolites, most strongly with riboflavin. These results suggest that some dissolved metabolites are linked to differing taxa across the transect. As the dominant Cyanobacteria genus shifts from *Prochlorococcus* in the south to *Synechococcus* in the north, dissolved pantothenic acid and PABA could be differentially derived from each genus, including in the northernmost samples (stations A4 and A7) where picoeukaryotes are also more abundant (Figure S3). In contrast, dissolved MTA and riboflavin were detected in relatively few samples in the south Atlantic, but were particularly abundant in the northernmost stations of the transect where picoeukaryotes and *Synechococcus* were more abundant than *Prochlorococcus* (Figures 6 and 7).

**Figure 6.**
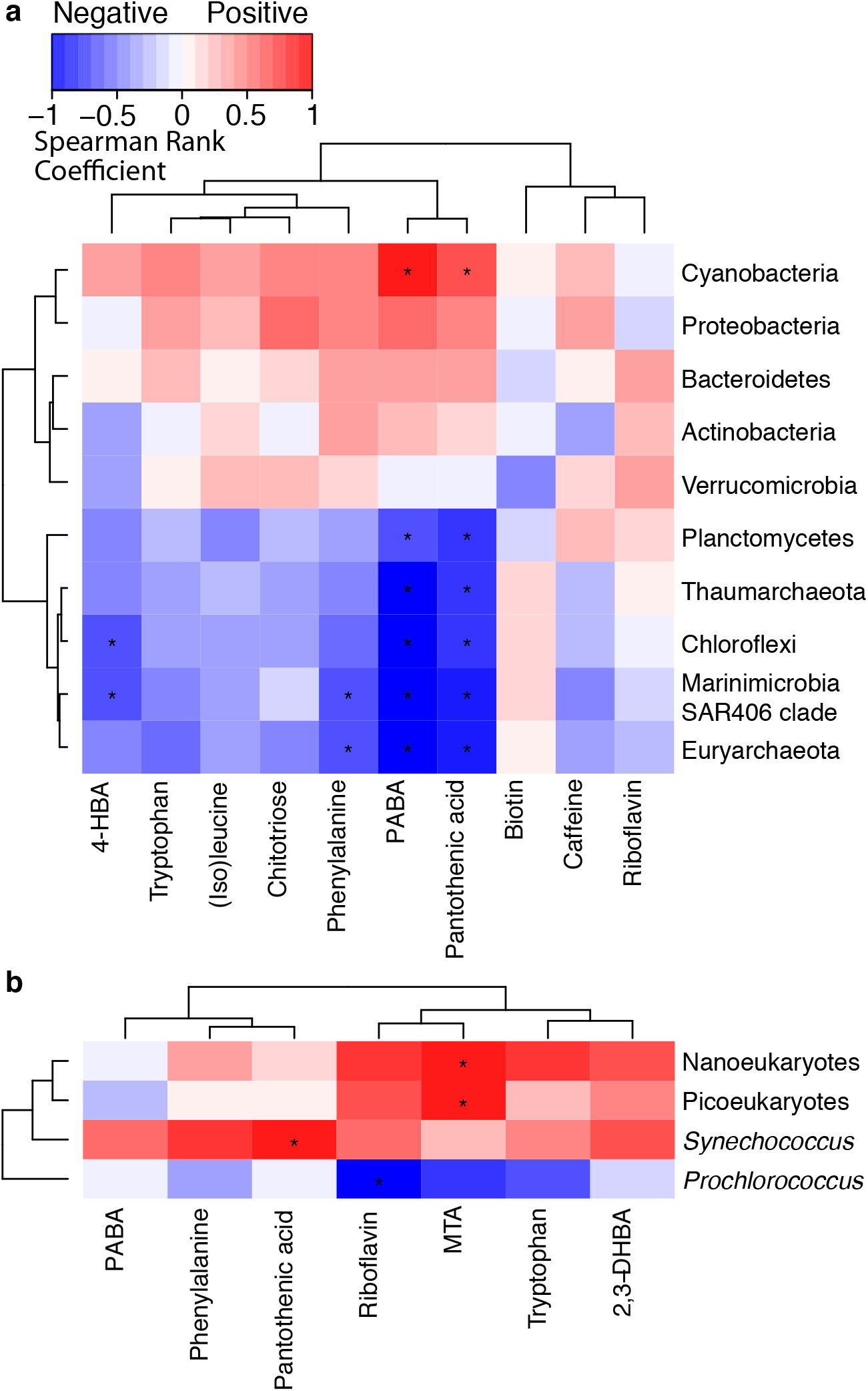
Spearman rank correlation coefficients plotted as a heatmap with hierarchical clustering showing the relationship between dissolved metabolites and abundance of microbial community members. Red indicates a positive correlation and blue indicates a negative correlation. * Indicates correlations with significant *p*-values. We include only samples collected in the upper 250 m as there were too few metabolites measured in the deeper samples (Table 2). (a) Comparison of metabolites to phyla from KN210-04 samples from 250 m and above (*p* < 0.05). (b) Comparison of AE1319 metabolite samples from the surface and DCM to groups identified with flow cytometry (*p* < 0.35).

**Figure 7.**
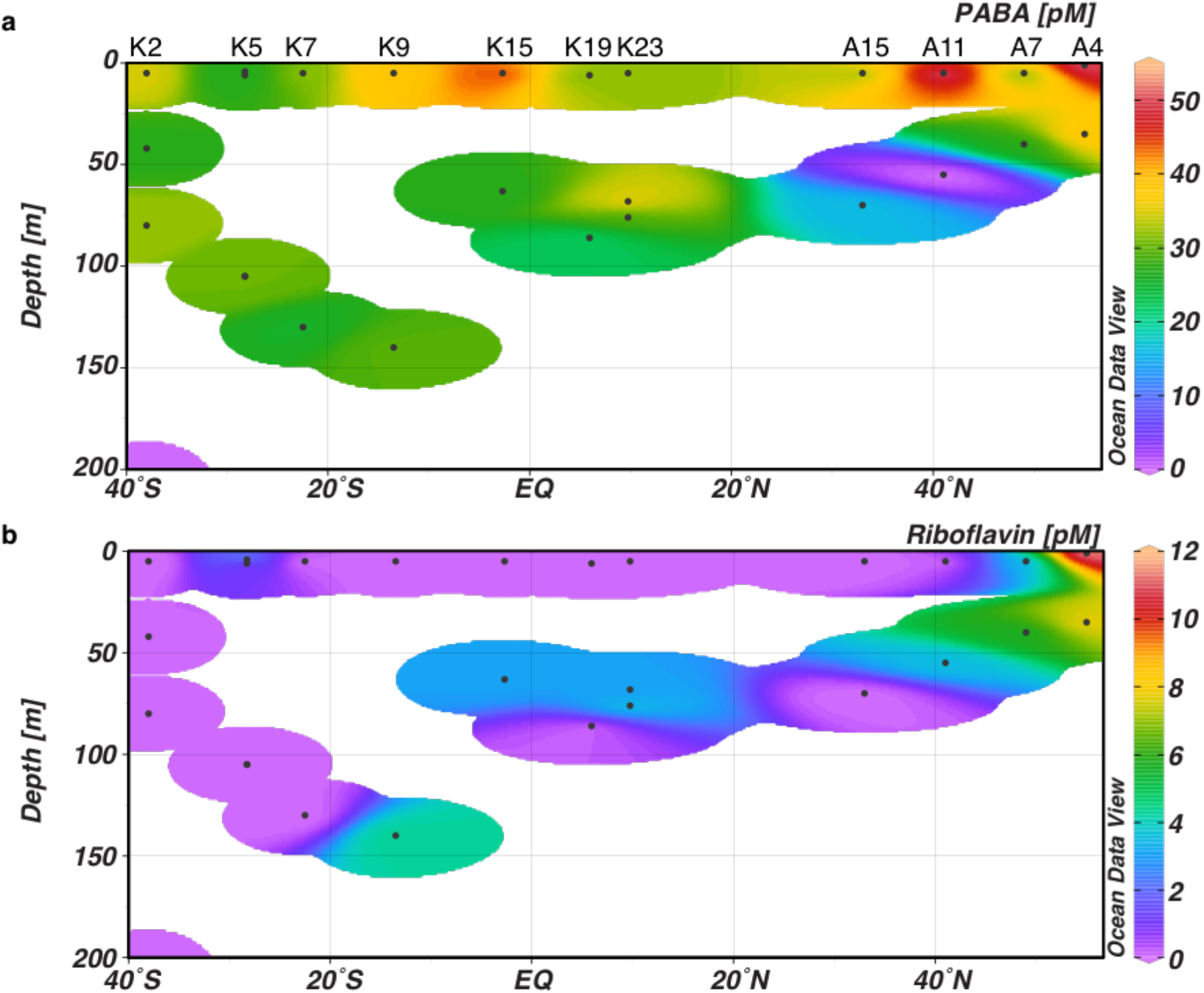
Concentrations of (a) dissolved 4-aminobenzoic acid (PABA) and (b) riboflavin (pM) in the top 200 m of the water column.

## Discussion

### Comparison to previously measured concentrations

Some of the metabolites presented here have been measured previously; in some cases, using similar mass spectrometry-based methods or with other analytical approaches. While the abundances of particulate metabolites are difficult to compare due to the diversity of ways in which molecules are normalized, concentrations of dissolved metabolites can be compared more readily with past values. Of the 18 dissolved metabolites quantified here, 12 have been measured previously in aquatic systems (Table S3). Two B vitamins, biotin (B_7_) and riboflavin (B_2_), had values of the same order of magnitude as the measurements made by Sañudo-Wilhelmy et al. (2012). A third B vitamin, thiamin (B_1_), had an order of magnitude higher concentration than that reported by Sañudo-Wilhelmy et al. (2012), but was only measured in one sample. Thiamin has a low extraction efficiency in the current protocol (2% in seawater; Johnson et al. 2017) and the resulting correction may introduce quantification error. When measured as dissolved free amino acids, (iso)leucine, phenylalanine, and tryptophan display a variety of concentrations ranging from hundreds to thousands of picomolar (Mopper and Lindroth 1982) in the upper 170 m of the Baltic Sea. In our study, concentrations varied from < 10 pM (deep ocean) to hundreds of pM (surface ocean). In short, dissolved metabolite concentrations obtained using this method agreed within an order of magnitude with existing data, where available.

### Many particulate metabolite distributions in the euphotic zone are linked to abundant microbial taxa

In the euphotic zone, many particulate metabolites have fairly similar profiles. Concentrations (Figure 3) differ most likely due to shifts in microbial community composition where members retain relatively constant intracellular concentrations of essential metabolites. For example, guanine (Figure 2) and AMP (Figure 4) were both elevated at the DCM and correlate with chlorophyll *a*. In the south and equatorial Atlantic, the phyla of Cyanobacteria, Actinobacteria, Proteobacteria, and Bacteroidetes positively correlated with particulate guanine, AMP, and other metabolites with similar profiles suggesting that they may be sources of these metabolites and thus have a strong influence on their distributions (Figure 3a). In the north Atlantic, nanoeukaryotes, picoeukaryotes, and *Synechococcus* were generally positively correlated with a similar set of metabolites, while *Prochlorococcus* was negatively correlated with these metabolite abundances (Figure 3b). This is likely a function of which taxa are the most abundant as these taxa will have the greatest impact on essential metabolite concentrations. It is also consistent with biovolume considerations, wherein larger microbes such as the pico- and nanoeukaryotes and *Synechococcus* carry larger metabolite concentrations due to their size, relative to the small *Prochlorococcus* (Kirchman 2008).

In contrast, some metabolites were not positively correlate with microbial abundance (Figure 3), including two B vitamins, a number of amino acids, and a nucleobase (adenine) and nucleobase derivatives (xanthine, uracil, caffeine). Adenine, a purine nucleobase, is an example of a metabolite that has weak or negative correlations with the dominant microbial taxa. It is measured in most surface samples, but in relatively few DCM samples compared to metabolites that positively correlate with Cyanobacteria, Proteobacteria and/or pico- and nanoeukaryotes. It is particularly striking that adenine was not detected at the Station K15 DCM where nutrients were being upwelled. In contrast, some of its highest concentrations were measured in the surface at Stations K5 and K7 in the oligotrophic South Atlantic Subtropical Gyre. This distinct pattern in particulate adenine distribution may be linked to how microbes respond to nutrient stress. In culture, *Escherichia coli* responds to both carbon and nitrogen starvation by elevating intracellular concentrations of adenine (Brauer et al. 2006). While the mechanism for this increase is not known, intracellular adenine concentrations play a regulatory role by binding to certain riboswitches (reviewed by Winkler and Breaker 2005). There are some exceptions to this trend of particulate adenine concentrations being elevated in oligotrophic samples. In particular, adenine was not detected at Station K9 in the gyre and the greatest adenine abundance was at Station K23 in the Amazon River plume. Thus, particulate adenine concentrations may be driven by something more complex than simply nutrient limitation, such as type or quality of nutrients, and variable species-specific responses. Metabolite distributions that are independent of biomass may highlight subtle and perhaps highly transient differences in metabolic state that are not reflected in more conservatively maintained intracellular metabolites.

### Variable patterns in the distribution of dissolved metabolites

Not all particulate and dissolved metabolites could be compared because some metabolites are not retained on the PPL polymer used to extract the dissolved metabolites (Johnson et al. 2017). In addition, some dissolved metabolites were not detected in particulate samples. Nevertheless, where data could be compared between particulate and dissolved pools, we observed greater variability in the distribution of dissolved metabolites. In rare cases, such as tryptophan and phenylalanine, the particulate and dissolved abundances appeared to co-vary (phenylalanine, r^2^ = 0.77; tryptophan, r^2^ = 0.73), implying a direct relationship between intracellular production and subsequent release into the water, as well as a fairly constant rate of removal. More frequently, we observed distinct distributions between the dissolved and particulate pools, and, in some cases, notable differences between the distributions of dissolved metabolites in the northern and southern portions of the transect (Figure 7; Figure 8).

**Figure 8.**
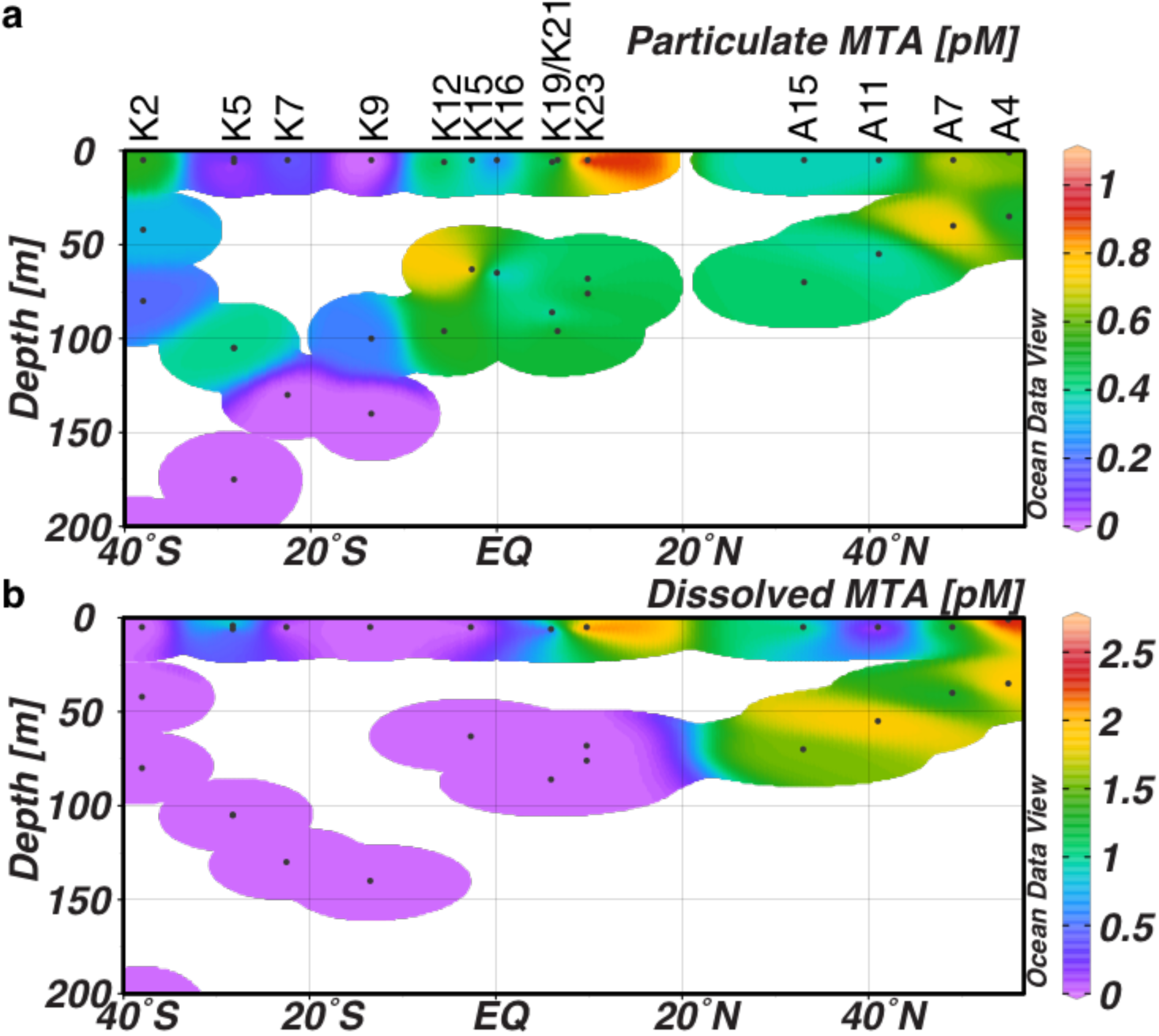
(a) Particulate 5-methylthioadenosine (MTA) normalized to the total volume filtered, and (b) pM dissolved MTA across the transect.

For example, PABA and pantothenic acid were positively correlated with Cyanobacteria in the southern portion of the transect and pantothenic acid was positively correlated with *Synechococcus* in the northern portion of the transect (Figure 6). Both PABA and pantothenic acid were particularly elevated in surface samples and measured throughout the transect (Figure 7a), suggesting a common source such as the Cyanobacteria. Due to its high rates of photosynthesis (Hartmann et al. 2014), *Prochlorococcus* can release between 9 and 24% of its initially assimilated inorganic carbon as DOC (Bertilsson et al. 2005). Of that, 4-20% can be low molecular weight carboxylic acids which include molecules like PABA and pantothenic acid (Bertilsson et al. 2005). Functionally, pantothenic acid is synthesized by plants and microbes and is a precursor to coenzyme A and acyl carrier protein. PABA is a precursor to the vitamin folate (Gibson and Pittard 1968). Both of these molecules could be valuable micronutrients within the microbial community. In a recent study identifying auxotrophy in the genomes of host-associated gram negative bacteria, some bacteria were predicted to be auxotrophic for both of these metabolites (Seif et al. 2020). Although there is no current evidence showing auxotrophy for these molecules in marine microbes, sustained environmental presence could provide the right conditions for loss of those biosynthetic pathways in some microbes (D’Souza and Kost 2016).

In contrast to PABA and pantothenic acid, riboflavin was more prevalent in the DCM and the northern portion of the transect (Figure 7b). Riboflavin has no correlation with any prokaryotic phyla in the south and equatorial Atlantic. However, in the north Atlantic, while the p-values are not statistically significant, we determined a weak positive correlation with *Synechococcus* and pico/nanoeukaryotes, and a significant negative correlation with *Prochlorococcus* (Figure 6). These correlations suggest that the source of dissolved riboflavin might be eukaryotes rather than Cyanobacteria. Riboflavin is a vitamin that could play a role as a micronutrient within the microbial community. Unlike some other B vitamins, auxotrophy for riboflavin has not been documented to our knowledge. However, it is likely that microbes would assimilate available riboflavin rather than synthesize it *de novo*. In non-marine systems, riboflavin plays other roles such as mimicking quorum sensing molecules (Rajamani et al. 2008) and priming plant defense responses (Zhang et al. 2009). The versatility of this molecule suggests we may not fully understand its role in marine systems.

Dissolved MTA had a similar distribution to riboflavin but was detected in fewer samples in the south Atlantic (Figure 8b). Nevertheless, it was detected in many particulate samples in both the north and south Atlantic, indicating an intracellular presence throughout the transect (Figure 8a). This is consistent with our understanding of MTA biochemistry, wherein intracellular excess MTA inhibits the enzymes that produce it, leading to relatively constant internal concentrations (Raina et al. 1982; Parsek et al. 1999). The latitudinal profile of particulate MTA shows a similar distribution to the other metabolites linked to the most common prokaryotic phyla along the transect. The highest abundances of particulate MTA are at stations A4, A7, the DCM at K15, and the surface at K23. The particulate MTA abundances were lower, but still elevated, in most DCM samples and in the north Atlantic where it was positively correlated with pico- and nano-eukaryotes (Figure 3; Figure 8a). In contrast, dissolved MTA was only measured in the north Atlantic and at the surface in K23 (Amazon River plume station; Figure 8b). This difference in MTA distributions between particulate and dissolved pools suggests that the intracellular concentration of MTA does not directly determine the dissolved MTA concentration, unlike the aromatic amino acids tryptophan and phenylalanine. In the south Atlantic, dissolved MTA was detected in too few samples to compare to the community composition. In the north Atlantic, like riboflavin, dissolved MTA was positively correlated with pico- and nanoeukaryotes (Figure 6), suggesting that in the dissolved phase MTA might be associated with eukaryotic taxa. Alternatively, while this cannot be determined from this dataset, there could be a more rapid removal process for dissolved MTA in the south Atlantic that maintains the dissolved concentrations below detection. However, a comparison of some representative genomes suggests that the distribution of dissolved MTA could be shaped by differences in a metabolic pathway between taxa. In culture experiments, the Cyanobacterium, *Synechococcus elongatus*, does not release MTA (Fiore et al. 2015) while the heterotrophic bacterium, *Ruegeria pomeroyi*, does (Johnson et al. 2016). According to the KEGG database, *S. elongatus,* and other *Synechococcus* and *Prochlorococcus* species lack the final gene in the methionine salvage pathway through which MTA is recycled back to methionine, providing the possibility that MTA could be further transformed through five additional enzymatic reactions before being released from the cell (Kanehisa and Goto 2000; Kanehisa et al. 2012). In contrast, *R. pomeroyi*, has a truncated methionine salvage pathway with only one additional possible downstream reaction after MTA. This reaction results in a phosphorylated metabolite that would be a costly waste product under low phosphorus conditions (Kanehisa and Goto 2000; Kanehisa et al. 2012). This suggests that rather than perform the final phosphorylation reaction, MTA might be released as a waste product as was observed in Johnson et al. (2016). Similarly, the picoeukaryotes *Ostreococcus tauri* and *Micromonas pusilla* also have a truncated methionine salvage pathway that ends with the same phosphorylated metabolite as *R. pomeroyi* (Kanehisa and Goto 2000; Kanehisa et al. 2012), suggesting that a higher abundance of picoeukaryotes could result in more dissolved MTA.

### Particulate metabolites that persist in the deep ocean

The metabolites measured in the deep ocean must either be derived from its unique resident microbial community or be delivered from the surface ocean by sinking particles. Interestingly, in a study that sought to establish a degradation index for particulate organic matter based on amino acid composition, Dauwe et al. (1999) found that threonine, arginine, aspartic acid and glycine increased as a mole percent of total amino acids with increasing organic matter degradation. While we did not quantify threonine, aspartic acid, or glycine in this study, we observed arginine in many deep samples (Figure 5a). While these amino acids are used as part of a degradation index for particulate organic matter (Dauwe et al. 1999), it is unclear whether their increased proportion in deep particulate organic matter is due to *in situ* production or selective preservation on sinking particles. We measured tryptophan and phenylalanine throughout the deep particulate samples. These amino acids are not linked to organic matter degradation according to Dauwe et al. (1999), suggesting that they might be produced by the microbial community at depth. However, data from sinking particles collected at 150 m along the south Atlantic transect indicate that phenylalanine and tryptophan are present in these sinking particles all along the transect, while arginine was only detected in ∼25% of the sinking particle samples (Johnson et al. 2020) suggesting that phenylalanine and tryptophan could be delivered by sinking particles but that arginine might be generated *in situ*.

Other particulate metabolites that we might expect to measure in the deep ocean include osmolytes, which are present at high concentrations in the cytosol of cells. However, glycine betaine and glutamic acid, both common osmolytes across all domains of life (Yancey et al. 1982) and present at high concentrations in the upper water column in this study were not detected in the deep ocean. By contrast, guanine and phenylalanine had similar concentrations to glycine betaine and glutamic acid in the euphotic zone, but were also measured in the deep ocean. This suggests that glycine betaine and glutamic acid decrease more quickly than biomass with depth. This may indicate that other molecules are used as osmolytes by deep-sea microbes due to differing requirements of organisms living in the colder, higher pressure waters of the ocean interior. Deep sea invertebrates have been shown to use different osmolytes than surface organisms but, to our knowledge, this has not been examined in marine microbes (Yancey et al. 2002). The amino acid proline also functions as an osmolyte (Burg and Ferraris 2008) and was measured in many deep samples, perhaps indicating a preferential use of this molecule as an osmolyte at high hydrostatic pressure and low temperatures (Figure 5b). In laboratory experiments, proline has been found to increase intracellularly with increasing osmolarity in *Bacillus subtilis* (Brill et al. 2011) and in *Sulfurimonas denitrificans* (Götz et al. 2018). While *S. denitrificans* was isolated from a tidal mudflat, other species in this genus have been found around hydrothermal vents in the deep ocean (Götz et al. 2018). It has not been widely characterized in studies of marine amino acid distributions, but recent work found that L-proline comprised 46% of the total (free and hydrolysable) amino acids in bacterial-sized particles (0.2 µm - 1.2 µm; Takasu and Nagata 2015). These studies support the idea that proline may be a substantial component of the cytosol of some cells in marine environments, including in the deep ocean. In the surface ocean osmolytes like DMSP are important sources of organic carbon that support heterotrophic growth, and it is possible that molecules such as proline may serve a similar function for deep ocean heterotrophs.

## Conclusion

By simultaneously measuring a suite of metabolites in a variety of oceanic regions and depths, we have developed a better understanding of metabolite variability in the ocean. We found that a group of metabolites remains constant relative to biomass, specifically associated with Cyanobacteria, Proteobacteria and/or eukaryote abundances, while other metabolites exhibited distinct distributions. Metabolites that deviated from biomass include nucleic acids, B vitamins, amino acids, and a variety of metabolic intermediates. While in many cases these metabolites have been studied in a biomedical context or in a model organism and their function in the cell is known, surprisingly little is known about how these molecules respond to environmental cues or their specificity within certain species (particularly in the ocean). This creates challenges in interpreting these data in the environment. There are surprises, such as the finding that particulate adenine was distributed in way that is not linked to abundant prokaryotic taxa, that dissolved metabolites like PABA and pantothenic acid might be linked to Cyanobacterial abundance while MTA and riboflavin appeared linked to pico- and nanoeukaryotes, and that proline was a prevalent amino acid in the deep sea. This dataset provides a context for future marine metabolomics work to constrain the ways in which specific metabolites may act as currencies that link diverse groups of microbes. Integrating metabolomics datasets with other omics datasets in the context of environmental parameters will allow further testing of hypotheses proposed here. In addition, controlled culture experiments can continue to expand our understanding of how metabolites respond to the environment, and additional field studies will complete the picture of metabolite distributions in other ocean regions. As this understanding is expanded, the mechanisms that control the flux of these metabolites through the marine food web and carbon cycle may be clarified and more accurately predicted.

## Acknowledgments

Funding for this work came from a National Science Foundation grant (Grant OCE-1154320 to EBK and KL). The instruments in the WHOI FT-MS Facility were purchased with support from the GBMF and NSF. Support for WMJ was provided by a National Defense Science and Engineering Fellowship. We thank the captains and crew of the R/V *Knorr* and the R/V *Atlantic Explorer*, Catherine Carmichael, Gwenn Hennon (Chl *a* calibration on KN210-04), and Erin Eggleston (prokaryotic cell counts on KN210-04) for helping to make this dataset possible. Targeted metabolomics datasets have been archived with MetaboLights (study # MTBLS1752). Sequencing was performed under the auspices of the US Department of Energy (DOE) JGI Community Science Program (CSP) project (CSP 1685) supported by the Office of Science of US DOE Contract DE-AC02-05CH11231. Additional work related to sample collection and processing was supported by the G. Unger Vetlesen and Ambrose Monell Foundations, the Natural Sciences and Engineering Research Council of Canada (NSERC), the Canadian Institute for Advanced Study (CIFAR) and the Canada Foundation for Innovation through grants awarded to SJH. MPB was supported by a CIFAR Global Scholarship and NSERC postdoctoral fellowship.

## Notes

### Competing Interest Statement

The authors have declared no competing interest.

## References

Aluwihare, L., D. Repeta, and R. Chen. 1997. A major biopolymeric component to dissolved organic carbon in surface sea water. Nature 387: 166–169.

Ankrah, N. Y. D., A. L. May, J. L. Middleton, and others. 2014. Phage infection of an environmentally relevant marine bacterium alters host metabolism and lysate composition. ISME J. 8: 1089–1100. doi:10.1038/ismej.2013.216

Arar, E. J., and G. B. Collins. 1997. Method 445.0: In vitro determination of chlorophyll a and pheophytin a in marine and freshwater algae by fluorescence.

Barak-Gavish, N., M. J. Frada, P. A. Lee, and others. 2018. Bacterial virulence against an oceanic bloom-forming phytoplankter is mediated by algal DMSP. 1–38.

Bertilsson, S., O. Berglund, M. J. Pullin, and S. W. Chisholm. 2005. Release of dissolved organic matter by Prochlorococcus. Vie Milieu 55: 225–231. doi:10.4319/lo.2007.52.2.0798

Brauer, M. J., J. Yuan, B. D. Bennett, W. Lu, E. Kimball, D. Botstein, and J. D. Rabinowitz. 2006. Conservation of the metabolomic response to starvation across two divergent microbes. Proc. Natl. Acad. Sci. U. S. A. 103: 19302–19307. doi:10.1073/pnas.0609508103

Brill, J., T. Hoffmann, M. Bleisteiner, and E. Bremer. 2011. Osmotically controlled synthesis of the compatible solute proline is critical for cellular defense of *Bacillus subtilis* against high osmolarity. J. Bacteriol. 193: 5335–5346. doi:10.1128/JB.05490-11

Burg, M. B., and J. D. Ferraris. 2008. Intracellular organic osmolytes: Function and regulation. J. Biol. Chem. 283: 7309–7313. doi:10.1074/jbc.R700042200

Carini, P., E. O. Campbell, J. Morré, and others. 2014. Discovery of a SAR11 growth requirement for thiamin’s pyrimidine precursor and its distribution in the Sargasso Sea. ISME J. 8: 1727–38. doi:10.1038/ismej.2014.61

Carlson, C. A., and D. A. Hansell. 2015. DOM sources, sinks, reactivity, and budgets, p. 65–102. In D.A. Hansell and C.A. Carlson [eds.], Biogeochemistry of marine dissolved organic matter. Elsevier.

Chambers, M. C., B. Maclean, R. Burke, and others. 2012. A cross-platform toolkit for mass spectrometry and proteomics. Nat. Biotechnol. 30: 918–920. doi:10.1038/nbt.2377

Clasquin, M. F., E. Melamud, and J. D. Rabinowitz. 2012. LC-MS data processing with MAVEN: A metabolomic analysis and visualization engine. Curr. Protoc. Bioinforma. 37: 1–23. doi:10.1002/0471250953.bi1411s37

Croft, M. T., M. J. Warren, and A. G. Smith. 2006. Algae need their vitamins. Eukaryot. Cell 5: 1175–1183. doi:10.1128/EC.00097-06

D’Souza, G., and C. Kost. 2016. Experimental evolution of metabolic dependency in bacteria. PLOS Genet. 12: 1–27. doi:10.5061/dryad.9b47j

Dauwe, B., J. J. Middelburg, P. M. J. Herman, and C. H. R. Heip. 1999. Linking diagenetic alteration of amino acids and bulk organic matter reactivity. Limnol. Oceanogr. 44: 1809–1814. doi:10.4319/lo.1999.44.7.1809

Dittmar, T., B. Koch, N. Hertkorn, and G. Kattner. 2008. A simple and efficient method for the solid-phase extraction of dissolved organic matter (SPE-DOM) from seawater. Limnol. Oceanogr. Methods 6: 230–235. doi:10.4319/lom.2008.6.230

Dittmar, T., and B. P. Koch. 2006. Thermogenic organic matter dissolved in the abyssal ocean. Mar. Chem. 102: 208–217. doi:10.1016/j.marchem.2006.04.003

Ducklow, H. W. 1999. Minireview: The bacterial content of the oceanic euphotic zone. FEMS Microbiol. 30: 1–10. doi:10.1016/S0168-6496(99)00031-8

Fiehn, O. 2002. Metabolomics–the link between genotypes and phenotypes. Plant Mol. Biol. 48: 155–71.

Fiore, C. L., K. Longnecker, M. C. Kido Soule, and E. B. Kujawinski. 2015. Release of ecologically relevant metabolites by the cyanobacterium, *Synechococcus elongatus* CCMP 1631. Environ. Microbiol. 17: 3949–3963. doi:10.1111/1462-2920.12899

Flombaum, P., J. L. Gallegos, R. A. Gordillo, and others. 2013. Present and future global distributions of the marine Cyanobacteria Prochlorococcus and Synechococcus. Proc. Natl. Acad. Sci. U. S. A. 110: 9824–9829. doi:10.1073/pnas.1307701110

Frias-Lopez, J., Y. Shi, G. W. Tyson, M. L. Coleman, S. C. Schuster, S. W. Chisholm, and E. F. Delong. 2008. Microbial community gene expression in ocean surface waters. Proc. Natl. Acad. Sci. U. S. A. 105: 3805–10. doi:10.1073/pnas.0708897105

Gebser, B., and G. Pohnert. 2013. Synchronized regulation of different zwitterionic metabolites in the osmoadaption of phytoplankton. Mar. Drugs 11: 2168–2182. doi:10.3390/md11062168

Gibson, F., and J. Pittard. 1968. Pathways of biosynthesis of aromatic amino acids and vitamins and their control in microorganisms. Bacteriol. Rev. 32: 465–492.

Götz, F., K. Longnecker, M. C. Kido Soule, K. W. Becker, J. Mcnichol, E. B. Kujawinski, and S. M. Sievert. 2018. Targeted metabolomics reveals proline as a major osmolyte in the chemolithoautotroph *Sulfurimonas denitrificans*. Microbiologyopen 1–7. doi:10.1002/mbo3.586

Hansell, D. A., C. A. Carlson, D. J. Repeta, R. Schlitzer, and I. N. Insights. 2009. Dissolved organic matter in the ocean. Oceanography 22: 202–211.

Hertkorn, N., R. Benner, M. Frommberger, P. Schmitt-Kopplin, M. Witt, K. Kaiser, A. Kettrup, and J. I. Hedges. 2006. Characterization of a major refractory component of marine dissolved organic matter. Geochim. Cosmochim. Acta 70: 2990–3010. doi:10.1016/j.gca.2006.03.021

Howard, E. M., C. A. Durkin, G. M. M. Hennon, F. Ribalet, and R. H. R. Stanley. 2017. Biological production, export efficiency, and phytoplankton communities across 8000 km of the South Atlantic. Global Biogeochem. Cycles 31: 1066–1088. doi:10.1002/2016GB005488

Johnson, W. M., M. C. Kido Soule, and E. B. Kujawinski. 2016. Evidence for quorum sensing and differential metabolite production by a marine bacterium in response to DMSP. ISME J. 10: 2304–2316. doi:10.1038/ismej.2016.6

Johnson, W. M., M. C. Kido Soule, and E. B. Kujawinski. 2017. Extraction efficiency and quantification of dissolved metabolites in targeted marine metabolomics. Limnol. Oceanogr. Methods 15: 417–428. doi:10.1002/lom3.10181

Johnson, W. M., K. Longnecker, M. C. Kido Soule, W. A. Arnold, M. P. Bhatia, S. J. Hallam, B. A. S. Van Mooy, and E. B. Kujawinski. 2020. Metabolite composition of sinking particles differs from surface suspended particles across a latitudinal transect in the South Atlantic. Limnol. Oceanogr. 65: 111–127. doi:10.1002/lno.11255

Kaiser, K., and R. Benner. 2009. Biochemical composition and size distribution of organic matter at the Pacific and Atlantic time-series stations. Mar. Chem. 113: 63– 77. doi:10.1016/j.marchem.2008.12.004

Kanehisa, M., and S. Goto. 2000. KEGG: kyoto encyclopedia of genes and genomes. Nucleic Acids Res. 28: 27–30.

Kanehisa, M., S. Goto, Y. Sato, M. Furumichi, and M. Tanabe. 2012. KEGG for integration and interpretation of large-scale molecular data sets. Nucleic Acids Res. 40: D109–D114. doi:10.1093/nar/gkr988

Kido Soule, M. C., K. Longnecker, W. M. Johnson, and E. B. Kujawinski. 2015. Environmental metabolomics: Analytical strategies. Mar. Chem. 177: 374–387. doi:10.1016/j.marchem.2015.06.029

Kiene, R. P., L. J. Linn, and J. a. Bruton. 2000. New and important roles for DMSP in marine microbial communities. J. Sea Res. 43: 209–224. doi:10.1016/S1385-1101(00)00023-X

Kirchman, D. L. 2008. Introduction and overview, p. 1–26. In D.L. Kirchman [ed.], Microbial Ecology of the Oceans. Wiley-Blackwell.

Kruskopf, M., and K. J. Flynn. 2006. Chlorophyll content and fluorescence responses cannot be used to gauge reliably phytoplankton biomass, nutrient status or growth rate. New Phytol. 169: 525–536. doi:10.1111/j.1469-8137.2005.01601.x

Kujawinski, E. B. 2011. The impact of microbial metabolism on marine dissolved organic matter. Ann. Rev. Mar. Sci. 3: 567–99. doi:10.1146/annurev-marine-120308-081003

Kujawinski, E. B., K. Longnecker, H. Alexander, S. T. Dyhrman, C. L. Fiore, S. T. Haley, and W. M. Johnson. 2017. Phosphorus availability regulates intracellular nucleotides in marine eukaryotic phytoplankton. Limnol. Oceanogr. Lett. 2: 119– 129. doi:10.1002/lol2.10043

Kujawinski, E. B., R. Del Vecchio, N. V. Blough, G. C. Klein, and A. G. Marshall. 2004. Probing molecular-level transformations of dissolved organic matter: Insights on photochemical degradation and protozoan modification of DOM from electrospray ionization Fourier transform ion cyclotron resonance mass spectrometry. Mar. Chem. 92: 23–37. doi:10.1016/j.marchem.2004.06.038

Lesser, M. P. 2006. Oxidative stress in marine environments: Biochemistry and physiological ecology. Annu. Rev. Physiol. 68: 253–278. doi:10.1146/annurev.physiol.68.040104.110001

Li, W. K. W., and W. G. Harrison. 2001. Chlorophyll, bacteria and picophytoplankton in ecological provinces of the North Atlantic. Deep. Res. Part II Top. Stud. Oceanogr. 48: 2271–2293. doi:10.1016/S0967-0645(00)00180-6

Lomas, M. W., J. A. Bonachela, S. A. Levin, and A. C. Martiny. 2014. Impact of ocean phytoplankton diversity on phosphate uptake. Proc. Natl. Acad. Sci. U. S. A. 111: 17540–17545. doi:10.1073/pnas.1420760111

Longnecker, K., M. C. Kido Soule, and E. B. Kujawinski. 2015. Dissolved organic matter produced by *Thalassiosira pseudonana*. Mar. Chem. 168: 114–123. doi:10.1016/j.marchem.2014.11.003

McCarthy, M., J. Hedges, and R. Benner. 1996. Major biochemical composition of dissolved high molecular weight organic matter in seawater. Mar. Chem. 55: 281– 297. doi:10.1016/S0304-4203(96)00041-2

Melamud, E., L. Vastag, and J. D. Rabinowitz. 2010. Metabolomic analysis and visualization engine for LC-MS data. Anal. Chem. 82: 9818–9826.

Mittenhuber, G. 2001. Phylogenetic analyses and comparative genomics of vitamin B6 (pyridoxine) and pyridoxal phosphate biosynthesis pathways. J. Mol. Microbiol. Biotechnol. 3: 1–20.

Mopper, K., and P. Lindroth. 1982. Diel and depth variations in dissolved free amino acids and ammonium in the Baltic Sea determined by shipboard HPLC analysis. Limnol. Oceanogr. 27: 336–347. doi:10.4319/lo.1982.27.2.0336

Moran, M. A., E. B. Kujawinski, A. Stubbins, and others. 2016. Deciphering ocean carbon in a changing world. Proc. Natl. Acad. Sci. U. S. A. 113: 3143–3151. doi:10.1073/pnas.1514645113

Noble, R. T., and J. A. Fuhrman. 1998. Use of SYBR Green I for rapid epifluorescence counts of marine viruses and bacteria. Aquat. Microb. Ecol. 14: 113–118. doi:10.3354/ame014113

Parsek, M. R., D. L. Val, B. L. Hanzelka, J. E. Cronan, and E. P. Greenberg. 1999. Acyl homoserine-lactone quorum-sensing signal generation. Proc. Natl. Acad. Sci. U. S. A. 96: 4360–5. doi:10.1073/pnas.96.8.4360

Paul, C., M. A. Mausz, and G. Pohnert. 2012. A co-culturing/metabolomics approach to investigate chemically mediated interactions of planktonic organisms reveals influence of bacteria on diatom metabolism. Metabolomics 9: 349–359. doi:10.1007/s11306-012-0453-1

Pohnert, G. 2000. Wound-activated chemical defense in a unicelular planktonic algae. Angew. Chemie Int. Ed. 30: 4352–4354.

R Core Team. 2015. R: A language and environment for statistical computing.

Rabinowitz, J. D., and E. Kimball. 2007. Acidic acetonitrile for cellular metabolome extraction from *Escherichia coli*. Anal. Chem. 79: 6167–6173.

Raina, A., K. Tuomi, and R. L. Pajula. 1982. Inhibition of the synthesis of polyamines and macromolecules by 5’-methylthioadenosine and 5’-alkylthiotubercidins in BHK21 cells. Biochem. J. 204: 697–703.

Rajamani, S., W. D. Bauer, J. B. Robinson, and others. 2008. The vitamin riboflavin and its derivative lumichrome activate the LasR bacterial quorum-sensing receptor. Mol. plant-microbe Interact. 21: 1184–92. doi:10.1094/MPMI-21-9-1184

Rich, J., M. Gosselin, E. Sherr, B. Sherr, and D. L. Kirchman. 1997. High bacterial production, uptake and concentrations of dissolved organic matter in the Central Arctic Ocean. Deep. Res. Part II Top. Stud. Oceanogr. 44: 1645–1663. doi:10.1016/S0967-0645(97)00058-1

Sañudo-Wilhelmy, S. A., L. S. Cutter, R. Durazo, and others. 2012. Multiple B-vitamin depletion in large areas of the coastal ocean. Proc. Natl. Acad. Sci. U. S. A. 109: 14041–5. doi:10.1073/pnas.1208755109

Schlitzer, R. 2016. Ocean Data View.

Seif, Y., K. Sonal, Y. Hefner, A. Anand, and L. Yang. 2020. Metabolic and genetic basis for auxotrophies in Gram-negative species. Proc. Natl. Acad. Sci. doi:10.1073/pnas.1910499117

Seyedsayamdost, M. R., R. J. Case, R. Kolter, and J. Clardy. 2011. The Jekyll-and-Hyde chemistry of *Phaeobacter gallaeciensis*. Nat. Chem. 3: 331–335. doi:10.1038/nchem.1002

Søndergaard, M., P. J. le B. Williams, G. Cauvet, B. Riemann, C. Robinson, S. Terzic, E. M. S. Woodward, and J. Worm. 2000. Net accumulation and flux of dissolved organic carbon and dissolved organic nitrogen in marine plankton communities. Limnol. Oceanogr. 45: 1097–1111.

Storey, J. D. 2002. A direct approach to false discovery rates. J. R. Stat. Soc. B 64: 479– 498. doi:10.1073/pnas.81.19.5921

Storey, J. D. 2010. False Discovery Rates Multiple Hypothesis Testing. 1–7.

Storey, J. D., and R. Tibshirani. 2003. Statistical significance for genomewide studies. Proc. Natl. Acad. Sci. U. S. A. 100: 9440–9445. doi:10.1073/pnas.1530509100

Takasu, H., and T. Nagata. 2015. High proline content of bacteria-sized particles in the Western North Pacific and its potential as a new biogeochemical indicator of organic matter diagenesis. Front. Mar. Sci. 2: 110. doi:10.3389/fmars.2015.00110

Thompson, P. A., M. X. Guo, P. J. Harrison, and J. N. C. Whyte. 1992. Effects of variation in temperature. II. On the fatty-acid composition of eight species of marine phytoplankton. J. Phycol. 28: 488–497. doi:10.1111/j.0022-3646.1992.00488.x

Winkler, W. C., and R. R. Breaker. 2005. Regulation of Bacterial Gene Expression By Riboswitches. Annu. Rev. Microbiol. 59: 487–517. doi:doi:10.1146/annurev.micro.59.030804.121336

Yancey, P. H., W. R. Blake, and J. Conley. 2002. Unusual organic osmolytes in deep-sea animals: Adaptations to hydrostatic pressure and other perturbants. Comp. Biochem. Physiol. Part A 133: 667–676. doi:10.1016/S1095-6433(02)00182-4

Yancey, P. H., M. E. Clark, S. C. Hand, R. D. Bowlus, and G. N. Somero. 1982. Classes of intracellular osmolyte systems and their distributions living with water stress: Evolution of osmolyte systems. Science (80-.). 217: 1214–1222. doi:10.1126/science.7112124

Yentsch, C. S., and D. W. Menzel. 1963. A method for the determination of phytoplankton chlorophyll and phaeophytin by fluorescence. Deep Sea Res. Oceanogr. Abstr. 10: 221–231. doi:10.1016/0011-7471(63)90358-9

Zhang, S., X. Yang, M. Sun, F. Sun, S. Deng, and H. Dong. 2009. Riboflavin-induced priming for pathogen defense in Arabidopsis thaliana. J. Integr. Plant Biol. 51: 167– 174. doi:10.1111/j.1744-7909.2008.00763.x

Zubkov, M. V., M. A. Sleigh, G. A. Tarran, P. H. Burkill, and R. J. G. Leakey. 1998. Picoplanktonic community structure on an Atlantic transect from 50°N to 50°S. Deep. Res. Part I Oceanogr. Res. Pap. 45: 1339–1355. doi:10.1016/S0967-0637(98)00015-6

