## Supplemental Tables and Figures for "Insights into the controls on metabolite distributions along a latitudinal transect of the western Atlantic Ocean"

### Supplemental Information

**Text S1.** Comparison of particulate and dissolved samples to identify cell leakage.
**Table S1.** Mixed layer depths, euphotic zone depths, and DCM depths.
**Table S2.** Dilutions for each sample analyzed.
**Table S3.** Concentrations of dissolved metabolites measured in previous studies.
**Figure S1.** Comparison of prokaryotic cell abundance and total moles of particulate metabolites.
**Figure S2.** Temperature and salinity along the transect.
**Figure S3.** Abundances of nanoeukaryotes, picoeukaryotes, *Synechococcus*, and *Prochlorococcus* in the north Atlantic.
**Figure S4.** Abundance of prokaryotic phyla in the south and equatorial Atlantic.
**Figure S5.** Comparison of particulate and dissolved metabolite concentrations normalized to phenylalanine.

**Figure S6.** Distribution of total moles of targeted particulate metabolites.

**Text S1.**

We examined the possibility that dissolved metabolite concentrations could be affected by cell damage during filtration. Here, we compared the ratios of metabolites in the dissolved phase to the same ratios in the particulate phase. Our hypothesis was that with pervasive cell damage, bursting or leakage, the dissolved ratios would approach those of the particulate phase, assuming no preferential loss or change in these metabolites during this physical disruption. Below, we plot the ratio of particulate metabolite *x* / phenylalanine against the ratio of dissolved metabolite *x* / phenylalanine (Figure S5). We chose phenylalanine for this comparison because it was measured in most dissolved and particulate samples, thus providing a ubiquitous denominator. All the metabolites examined had samples where the metabolite was detected in only one of the two fractions, thus reducing the number of comparisons that could be made, but also supporting the contention that leakage was not likely to be a significant problem. Where a metabolite was detected in both fractions, some samples had similar particulate and dissolved ratios, in particular, riboflavin, 5-methylthioadenosine (MTA), and inosine. However, this was highly variable across the different metabolites and samples suggesting that there was no systematic release of these metabolites into the dissolved phase, as would be expected from cell leakage or physical lysis. We conclude that cell leakage, or breakage during filtration, had a minimal impact on our data.

**Table S1.** Mixed layer depth, euphotic zone depth, and deep chlorophyll maximum (DCM) depth along the transect. --- indicates stations where data was not available to determine the euphotic zone depth.

| Station | Mixed layer depth (m) | Euphotic zone depth (m) | DCM (m) |
|---|---|---|---|
| K2 | 98 | 129 | 80 |
| K5 | 70 | 128 | 105 |
| K7 | 54 | 117 | 130 |
| K9 | 48 | 135 | 140 |
| K12 | 68 | 102 | 96 |
| K15 | 48 | 83 | 63 |
| K16 | 50 | --- | 65 |
| K19 | 91 | 85 | 86 |
| K21 | 138 | 91 | 96 |
| K23 | 82 | 82 | 68 |
| A15 | 29 | 79 | 70 |
| A11 | 19 | --- | 55 |
| A7 | 33 | 86 | 40 |
| A4 | 31 | 42 | 35 |

**Table S2.** Dissolved sample dilutions. Due to ion suppression in the middle of the chromatogram certain samples were diluted, mostly samples below the euphotic zone.

| Station | Depth | Dilution |
|---|---|---|
| K2 | 5 | no dilution |
| K2 | 42 | no dilution |
| K2 | 80 | no dilution |
| K2 | 220 | no dilution |
| K2 | 754 | 2-fold |
| K2 | 2502 | 5-fold |
| K2 | 5110 | 2-fold |
| K7 | 5 | no dilution |
| K7 | 130 | no dilution |
| K7 | 250 | no dilution |
| K7 | 753 | 5-fold |
| K7 | 2500 | 2-fold |
| K7 | 4512 | 5-fold |
| K15 | 5 | no dilution |
| K15 | 63 | no dilution |
| K15 | 250 | no dilution |
| K15 | 750 | 2-fold |
| K15 | 2500 | no dilution |
| K15 | 5009 | no dilution |
| K23 | 5 | no dilution |
| K23 | 76 | no dilution |
| K23 | 68 | no dilution |
| K23 | 251 | no dilution |
| K23 | 248 | no dilution |
| K23 | 562 | 5-fold |
| K23 | 930 | 2-fold |
| K23 | 2500 | 2-fold |
| K23 | 3770 | 2-fold |
| K5 | 4 | no dilution |
| K5 | 6 | no dilution |
| K5 | 105 | no dilution |
| K5 | 105 | no dilution |
| K5 | 248 | no dilution |
| K9 | 5 | no dilution |
| K9 | 140 | no dilution |
| K9 | 253 | no dilution |
| K19 | 6 | no dilution |
| K19 | 86 | no dilution |
| K19 | 252 | no dilution |
| K12 | 6 | no dilution |
| K12 | 96 | no dilution |

| | | |
|---|---|---|
| K16 | 5 | no dilution |
| K16 | 65 | no dilution |
| K21 | 5 | no dilution |
| K21 | 96 | no dilution |
| A4 | 1 | no dilution |
| A4 | 35 | no dilution |
| A4 | 1500 | 2-fold |
| A4 | 3000 | 2-fold |
| A7 | 5 | no dilution |
| A7 | 40 | no dilution |
| A11 | 5 | no dilution |
| A11 | 55 | no dilution |
| A11 | 350 | 2-fold |
| A15 | 5 | no dilution |
| A15 | 70 | no dilution |
| A15 | 315 | 2-fold |
| A15 | 800 | 2-fold |
| A15 | 1500 | 2-fold |
| A15 | 3000 | 2-fold |

**Table S3.** Previous measurements of dissolved metabolites in the ocean.

| Compound | Dissolved (pM) | Reference |
|---|---|---|
| 2,3-Dihydroxybenzoic acid | 1-9† | (Longnecker et al. 2018) |
| 4-Aminobenzoic acid | 1-58† | (Longnecker et al. 2018) |
| 4-Hydroxybenzoic acid | 10-53† | (Longnecker et al. 2018) |
| 5-Methylthioadenosine | Not found | |
| Biotin | 10-200* | (Sañudo-Wilhelmy et al. 2012) |
| Caffeine | No non-anthro∞ | |
| Chitotriose | Not found | |
| Desthiobiotin | 5-100† | (Longnecker et al. 2018) |
| Indole 3-acetic acid | 5-400• | (Amin et al. 2015) |
| Inosine | Not found | |
| (Iso)leucine | 790-8000§; 210-270† | (Mopper and Lindroth 1982; Longnecker et al. 2018) |
| *N*-Acetylglutamic acid | Not found | |
| NAD | Not found | |
| Pantothenic acid | 4-50† | (Longnecker et al. 2018) |
| Phenylalanine | 400-4000§; 55-300† | (Mopper and Lindroth 1982; Longnecker et al. 2018) |
| Riboflavin | 0.2-5*; 0.6-38† | (Sañudo-Wilhelmy et al. 2012; Longnecker et al. 2018) |
| Thiamin | 25-350* | (Sañudo-Wilhelmy et al. 2012) |
| Tryptophan | 300-1000§; 3-110† | (Mopper and Lindroth 1982; Longnecker et al. 2018) |

*-Coastal California (1-800 m)
§-Baltic Sea (1-170 m)
•-North Pacific Ocean (5-150 m)
†-Seawater near hydrothermal vents and/or vent fluid. These concentrations were measured using the same method as in this study.
∞-Caffeine has only been measured in coastal environments with known anthropogenic sources of the molecule.

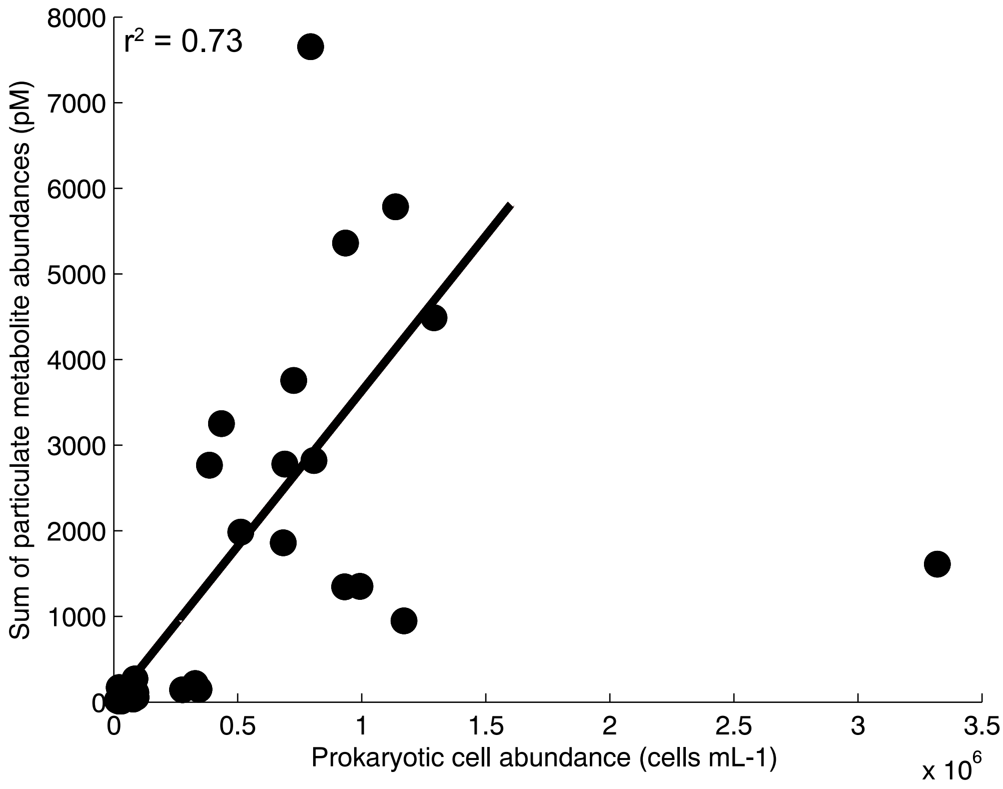


**Figure S1.** Cell counts plotted versus the total moles of targeted metabolites measured per liter of water filtered. Cell counts were performed on selected samples and show a linear correlation with the total moles of targeted metabolites measured in the particulate samples suggesting that this is a reasonable way to correct for biomass. One outlier has been discarded from the linear fit but is plotted in the right bottom corner. Pearson correlation test results: $r^2 = 0.73$, $p < 0.001$.

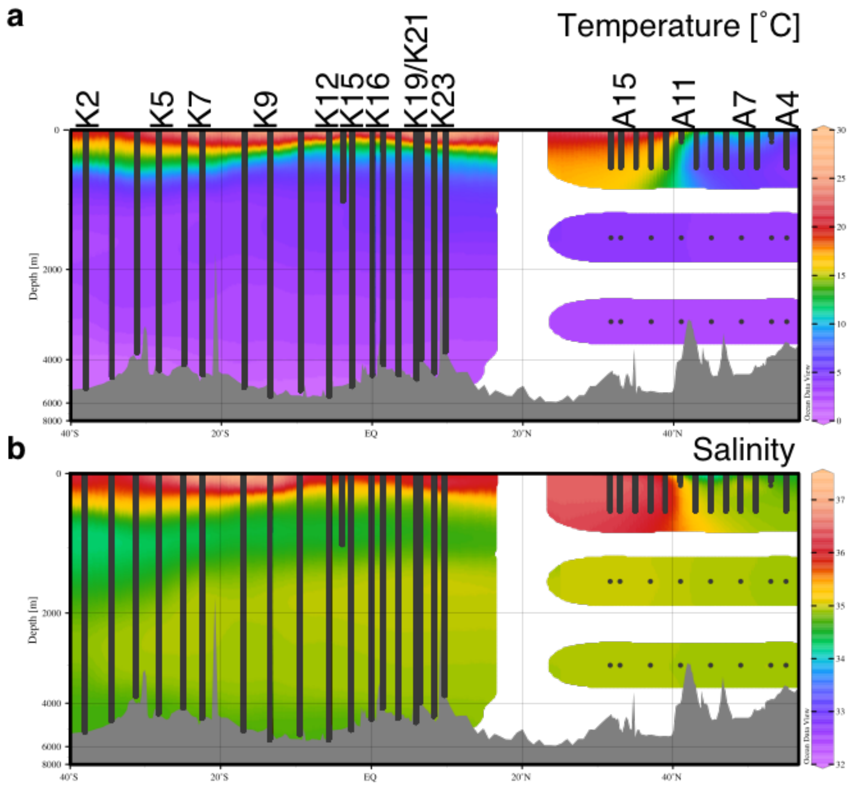


**Figure S2.** Temperature and salinity along the transect. Gray lines indicate CTD casts.

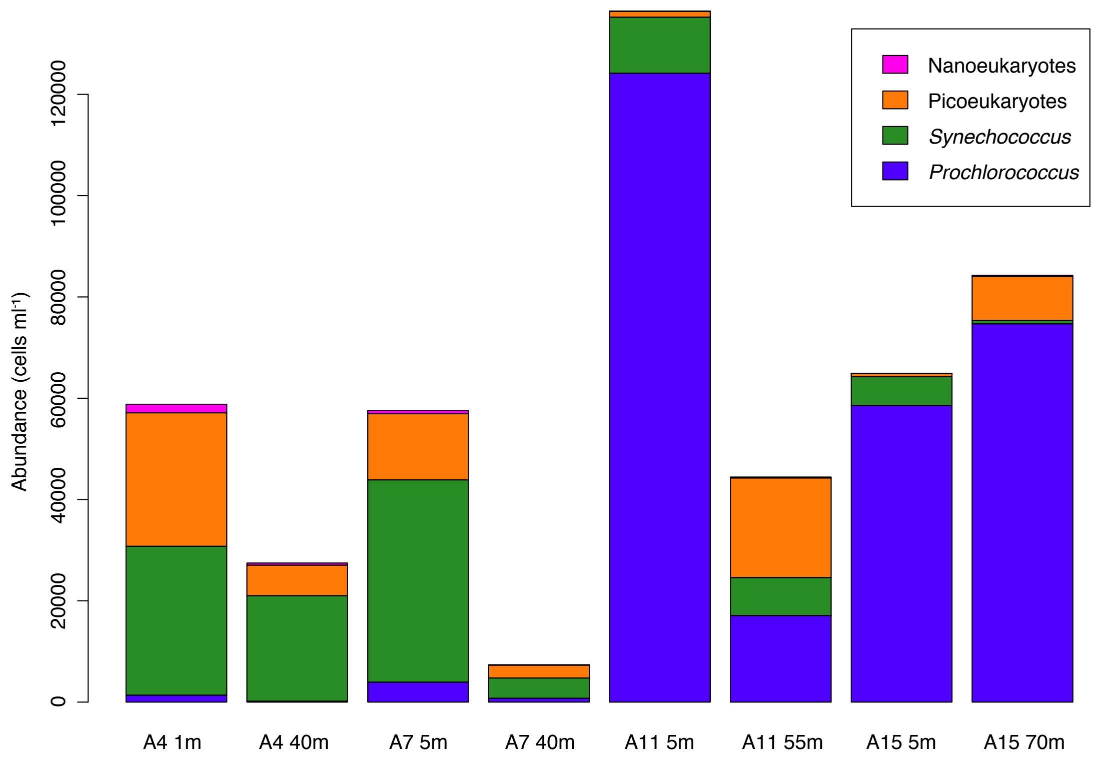


**Figure S3.** Abundance of nanoeukaryotes, picoeukaryotes, *Synechococcus*, and *Prochlorococcus* from cruise AE1319 in the north Atlantic. The numbers under each bar indicate the station number followed by sampling depth.

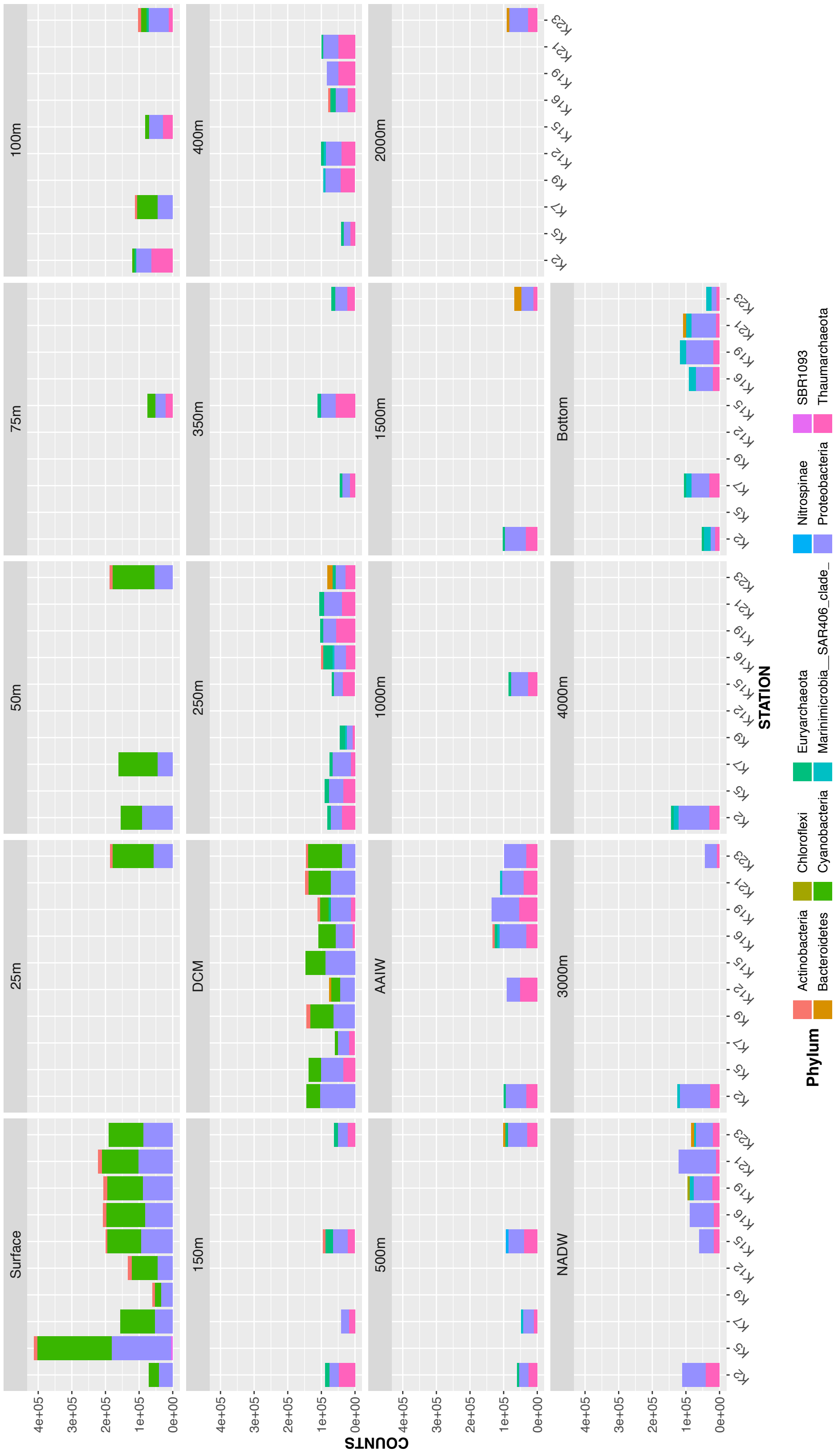


**Figure S4.** Abundance of prokaryotic phyla by counts of OTUs for cruise KN210-04 in the south Atlantic.

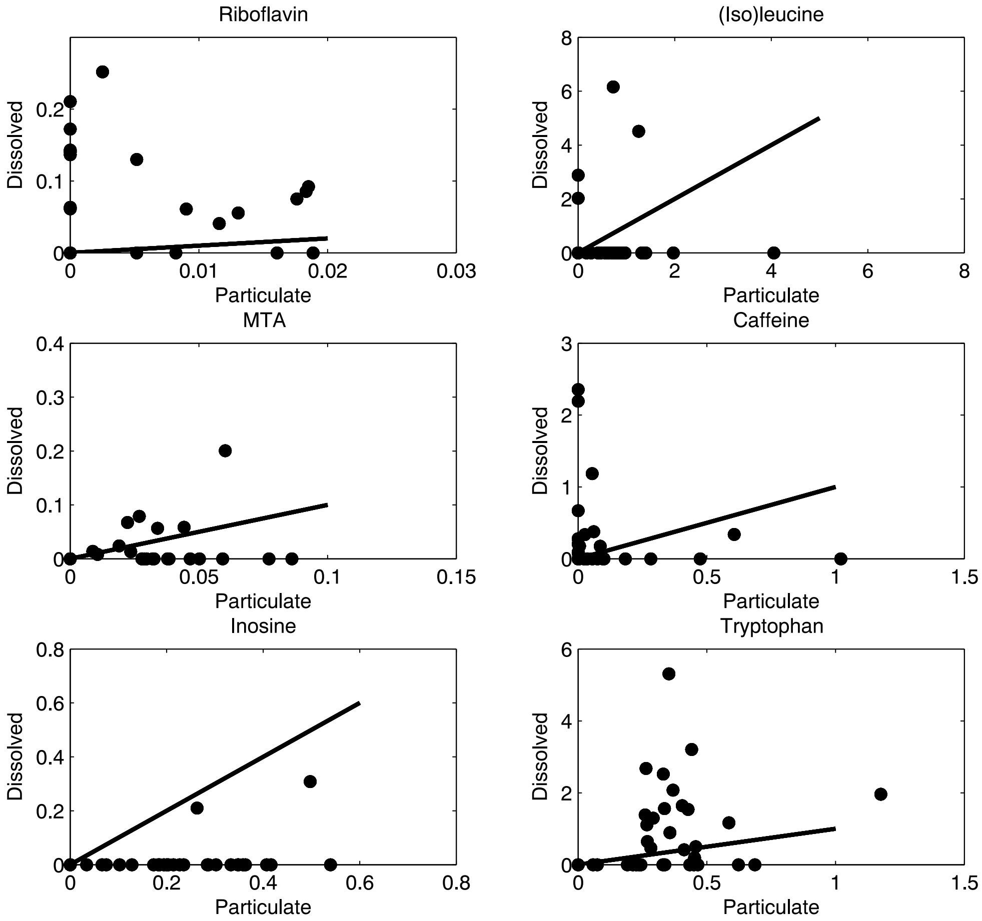


**Figure S5.** The particulate ratio of metabolite *x* / phenylalanine against the dissolved ratio of metabolite *x* / phenylalanine. Phenylalanine was measured in most dissolved and particulate samples. If metabolites were leaking from damaged cells during filtration, the ratio in the particulate phase should approach the ratio in the dissolved sample. The black line shows an equal relationship. This has been plotted for a) riboflavin, b) (iso)leucine, c) MTA, d) caffeine, e) inosine, f) tryptophan.

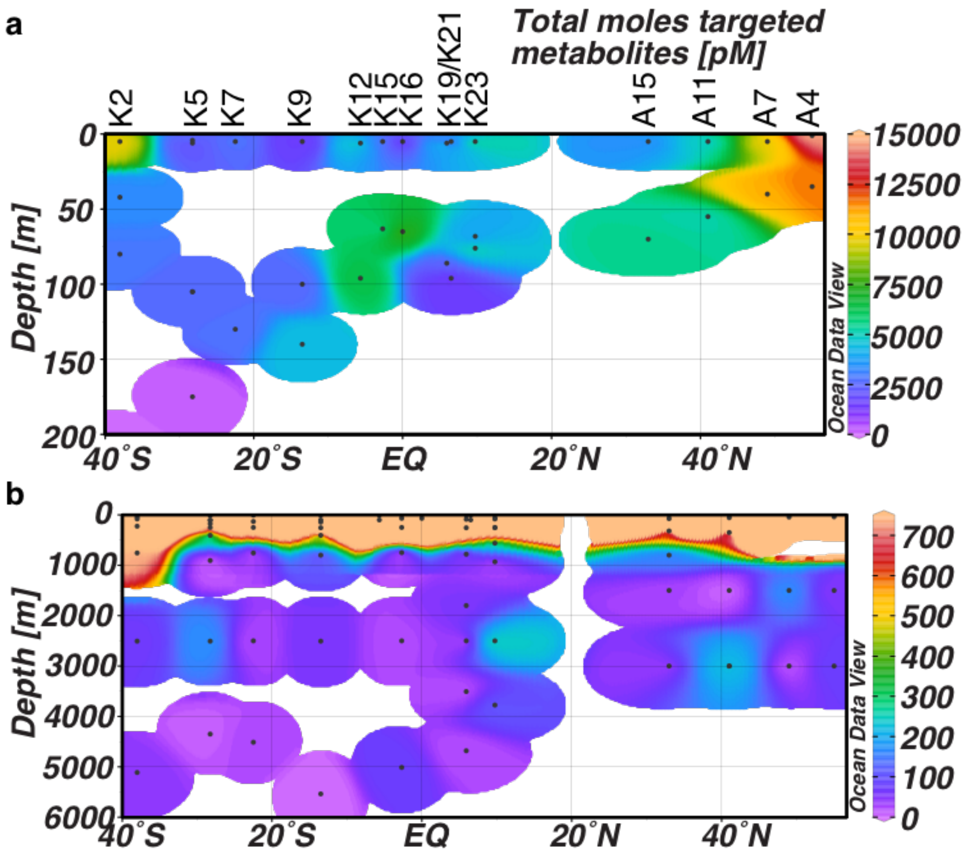


**Figure S6.** Distribution of total moles of particulate targeted metabolites over the transect, in the top 200 m (a) and at all depths (b).

**Supplemental References:**


Amin, S. A., L. R. Hmelo, H. M. van Tol, and others. 2015. Interaction and signalling between a cosmopolitan phytoplankton and associated bacteria. Nature **522**: 98–101. doi:10.1038/nature14488

Longnecker, K., S. M. Sievert, S. P. Sylva, J. S. Seewald, and E. B. Kujawinski. 2018. Dissolved organic carbon compounds in deep-sea hydrothermal vent fluids from the East Pacific Rise at 9°50′N. Org. Geochem. **125**: 41–49. doi:10.1016/j.orggeochem.2018.08.004

Mopper, K., and P. Lindroth. 1982. Diel and depth variations in dissolved free amino acids and ammonium in the Baltic Sea determined by shipboard HPLC analysis. Limnol. Oceanogr. **27**: 336–347. doi:10.4319/lo.1982.27.2.0336

Sañudo-Wilhelmy, S. A., L. S. Cutter, R. Durazo, and others. 2012. Multiple B-vitamin depletion in large areas of the coastal ocean. Proc. Natl. Acad. Sci. U. S. A. **109**: 14041–5. doi:10.1073/pnas.1208755109
